# MCM10 and SLD-2/RECQL4 jointly activate the CMG helicase during metazoan DNA replication initiation

**DOI:** 10.64898/2026.04.01.715773

**Authors:** Remi Sonneville, Cecile Evrin, Jane Wright, Yisui Xia, Karim P.M. Labib

## Abstract

Eukaryotic cells regulate the assembly and activation of the essential DNA helicase at the heart of the chromosome replication machinery, to ensure that the chromosomes are copied just once per cell cycle. The Mcm10 protein is essential for helicase activation in budding yeast, but an equivalent role for MCM10 orthologues in animal cells has not been explored. Moreover, complete deletion of the *mcm-10* gene is viable in the nematode *Caenorhabditis elegans*, suggesting the involvement of additional factors. Here we show that MCM-10 and a second factor called SLD-2 are recruited to chromatin after helicase assembly in the *C. elegans* early embryo and are jointly required for helicase activation. Moreover, deletion of the *Mcm10* gene is viable in mouse embryonic stem cells, but causes synthetic lethality in the absence of RECQL4, which is the orthologue of SLD-2 in vertebrate species. Helicase activation is blocked in the combined absence of MCM10 and RECQL4, mirroring the situation in *C. elegans*. These findings indicate that metazoan helicase activation requires two conserved factors that are mutated in human disease syndromes.

## Introduction

The proliferation of eukaryotic cells is dependent upon making a single copy of every chromosome during each cell cycle. The duplication of eukaryotic chromosomes is controlled by the regulated assembly and disassembly of the chromosome replication machinery, or replisome, to ensure that each chromosome is copied just once (Costa & Diffley, 2022). The replisome assembles around the 11-subunit DNA helicase known as CMG (<u>C</u>DC45, <u>M</u>CM2-7, <u>G</u>INS), which is built *in situ* at DNA replication origins in a two-step process that can only occur once per cell cycle. Once assembled, the nascent helicases are activated by a mechanism that remains poorly characterised in all species.

During the G1-phase of the cell cycle, two rings of the six MCM2-7 ATPases are loaded around double-stranded DNA (ds DNA) at replication origins, to form an MCM2-7 double-hexamer that lacks helicase activity (Frisbie & Bleichert, 2026; Stillman *et al*, 2025). Subsequently, when cells enter S-phase, the CDC45 protein and the 4-protein GINS complex are recruited to MCM2-7 double hexamers to form ‘head-to-head’ pairs of CMG helicase complexes, which initially still encircle ds DNA (Gambus *et al*, 2006; Lewis *et al*, 2022; Moyer *et al*, 2006).

Activation of the MCM2-7 ATPases within the double CMG complex leads to melting of ds DNA within the central cavity of the MCM2-7 hexamer, and also to extrusion of one of the two parental DNA strands through a ‘gate’ in the MCM2-7 ring (Costa & Diffley, 2022), likely between the MCM2 and MCM5 subunits (Weekes *et al*, 2026). This produces two active CMG helicases that encircle opposite DNA strands at a pair of nascent DNA replication forks. The two helicases translocate along DNA in a 3’ to 5’ direction (Fu *et al*, 2011), bypassing each other and then moving away from the origin in opposite directions (Douglas *et al*, 2018; Georgescu *et al*, 2017). At the same time, each CMG helicase associates with a set of partner proteins to form a replisome that commences DNA synthesis (Gambus *et al*., 2006; Gambus *et al*, 2009; Sengupta *et al*, 2013).

The loading of MCM2-7 double hexamers around origin DNA has been studied extensively, aided by biochemical reconstitution and structural analysis of intermediates (Costa & Diffley, 2022; Evrin *et al*, 2009; Frisbie & Bleichert, 2026; Remus *et al*, 2009; Stillman *et al*., 2025), principally in the budding yeast *Saccharomyces cerevisiae* but more recently with purified human proteins (Weissmann *et al*, 2024; Wells *et al*, 2025; Yang *et al*, 2024). These studies identified a conserved set of MCM2-7 loading factors and also revealed surprising mechanistic diversity, both within an individual species and also between yeast and humans.

The conversion of MCM2-7 double hexamers into a pair of CMG helicase complexes has also been reconstituted with purified budding yeast proteins (Yeeles *et al*, 2015), and intermediates in the CMG assembly reaction have been studied by cryo-electron microscopy (Costa & Diffley, 2022). Cdc45 is coupled to MCM2-7 double hexamers by Sld3 (Deegan *et al*, 2016; Li *et al*, 2025), whereas GINS is recruited by a combination of Sld2, Dpb11, and DNA polymerase epsilon (Pol ε) that subsequently synthesises the leading strand at DNA replication forks (Muramatsu *et al*, 2010). These factors are all conserved in diverse eukaryotic species. However, recent work indicates important mechanistic differences and additional complexity during CMG assembly in animal species, for which biochemical reconstitution of CMG assembly has yet to be achieved. Studies in the nematode *Caenorhabditis elegans* (Xia *et al*, 2023), the frog *Xenopus laevis* (Hashimoto *et al*, 2023; Kingsley *et al*, 2023; Lim *et al*, 2023), and also in mouse and humans (Evrin *et al*, 2023; Lim *et al*., 2023) showed that a protein called DONSON is essential for the recruitment of GINS to MCM2-7 double hexamers during CMG assembly. In humans, DONSON is mutated in a form of microcephalic primordial dwarfism (Reynolds *et al*, 2017; Schulz *et al*, 2018) that is typically caused by mutations in CMG components or assembly factors (Bellelli & Boulton, 2021; Klingseisen & Jackson, 2011; Nielsen-Dandoroff *et al*, 2023). However, DONSON appears to have been lost during fungal evolution and so is not present in yeast cells.

In budding yeast, Mcm10 is essential for the activation of nascent CMG helicases *in vivo* (van Deursen *et al*, 2012; Watase *et al*, 2012) and is sufficient for CMG activation during reconstituted replication reactions *in vitro* (Douglas *et al*., 2018). Similarly, Mcm10 is also required for CMG activation in the fission yeast, *Schizosaccharomyces pombe* (Kanke *et al*, 2012). However, studies of MCM10 orthologues in animal cells suggested the existence of other CMG activation factors. For example, depletion of more than 99% of MCM10 from *Xenopus laevis* egg extracts did not inhibit DNA replication *in vitro* (Chadha *et al*, 2016). Moreover, truncating mutations within *Drosophila* Mcm10 did not prevent the development of adult flies, though the resulting females had greatly reduced fertility (Reubens *et al*, 2015). Recently, worms lacking the first 96% of the coding sequence of *C. elegans mcm-10* were found to be viable and fertile (Xia *et al*., 2023), strongly indicating the existence of an MCM-10-independent pathway for CMG helicase activation in animal cells. Here we take a multi-dimensional approach based on *C. elegans* early embryos and mammalian cells, to show that the orthologues of yeast Sld2 and Mcm10 are jointly required for helicase activation but are dispensable for assembly of the CMG complex, during the initiation of chromosome replication in metazoa.

## Results

### CMG activation is delayed in the absence of MCM-10 or SLD-2 in the *C. elegans* early embryo

The first cell cycle of the *C. elegans* embryo provides a model system for studying the regulation of the metazoan CMG helicase *in vivo* (Sonneville *et al*, 2019; Sonneville *et al*, 2017; Sonneville *et al*, 2012; Xia *et al*, 2021; Xia *et al*., 2023). After fertilisation, DNA replication initiates in the female and male pronuclei on chromosomes that are still condensed after the second meiotic cell division (Figure EV1A). The chromosomes then decondense rapidly during early S-phase, in response to DNA unwinding by the newly activated CMG helicase (Sonneville *et al*, 2015). Therefore, chromosome decondensation is delayed when CMG assembly is blocked by depletion of CMG components (Sonneville *et al*., 2015), or by depletion of CMG assembly factors such as DNSN-1 that is the worm orthologue of mammalian DONSON (Xia *et al*., 2023).

In *mcm-10Δ* worms, chromosome decondensation was delayed during S-phase of the first embryonic cell cycle (Figure 1A, mCherry-H2B). Moreover, the GINS subunit PSF-1 was detected on condensed chromosomes throughout the period of the decondensation delay (Figure 1A, *mcm-10Δ*; Figure EV1B). Since the association of GINS with chromatin requires the prior recruitment of CDC-45 to double hexamers of MCM-2-7 (Xia *et al*., 2023), these data indicate that the CMG helicase assembles in *mcm-10Δ* worms but then remains inactive for an extended period, thereby delaying the initiation of DNA replication and chromosome decondensation. Correspondingly, DNA synthesis was found to be reduced at the start of the first cell cycle in *mcm-10Δ* compared to control, when embryos were treated with a short pulse of the nucleoside precursor 5-ethynyl-2’-deoxyuridine or EdU (Figure 1B-C). Subsequently, chromosome replication is completed and mitosis proceeds normally (Figure EV2A-C, *mcm-10Δ*), and *mcm-10Δ* embryos produce viable adult worms (Xia *et al*., 2023), suggesting that other factors are subsequently able to activate the CMG helicase, despite the absence of MCM-10.

**Figure 1.**
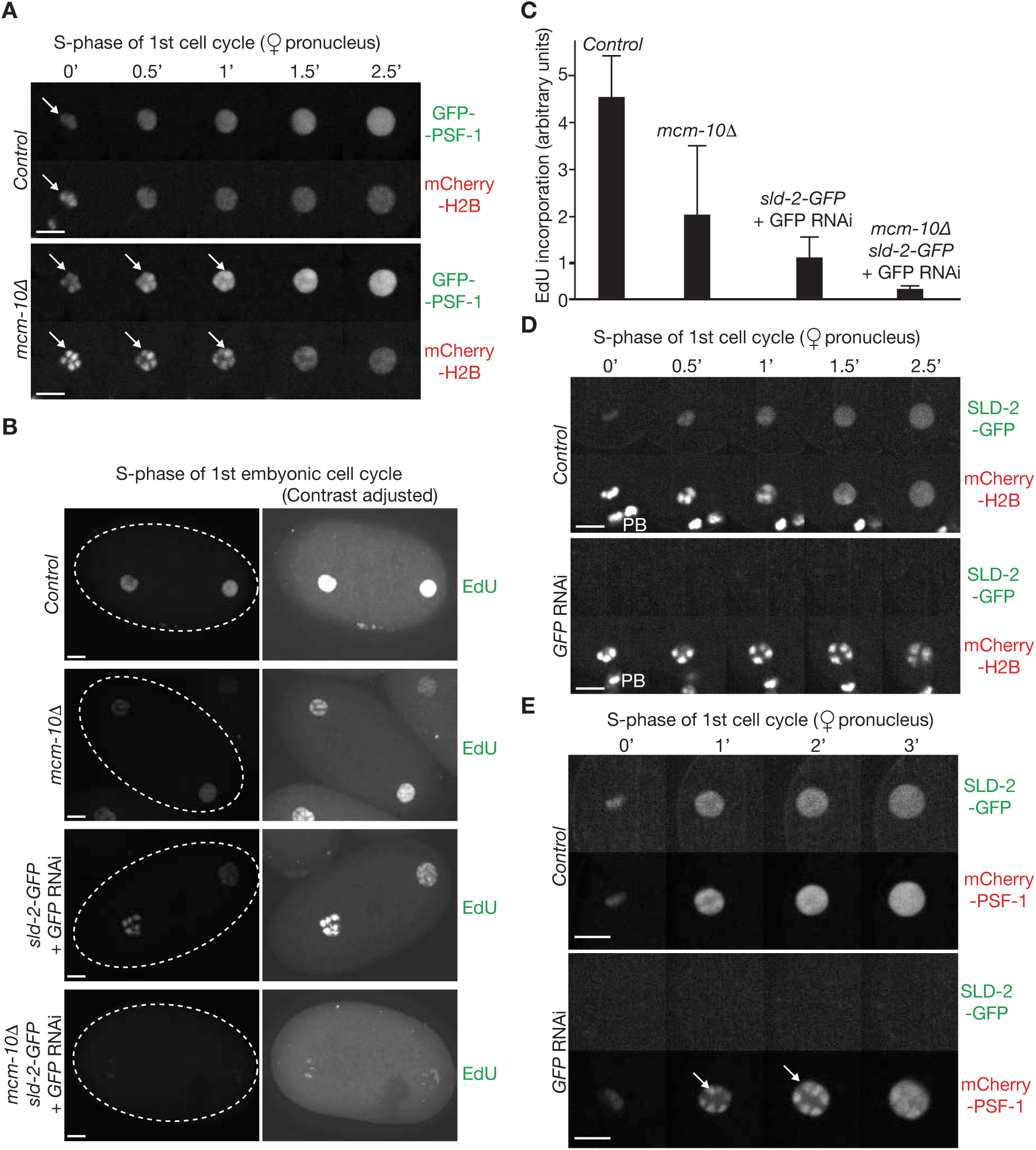
DNA replication is impaired after CMG helicase assembly in the absence of MCM-10 or SLD-2 in the *C. elegans* early embryo. (**A**) Chromosome decondensation was monitored for the indicated strains, upon entry into the first cell cycle of the *C. elegans* embryo. A sequence of images is presented, corresponding to progression of the female pronucleus through early S-phase at the indicated times. Arrows denote the association of GFP-PSF-1 with condensed chromosomes, which persist for longer in the absence of MCM-10. (**B**) Embryos of the indicated genotypes, treated as indicated with GFP RNAi, were given a 10-minute pulse of 20 mM EdU at the beginning of the first cell cycle. EdU was then detected as described in Materials and Methods. The right panels higher contrast versions of the images in the left panels. (**C**) EdU incorporation from (B) was quantified as described in Materials and Methods. The data show the mean and standard deviation from three independent experiments. (**D**) Worms expressing SLD-2-GFP and mCherry-Histone H2B were treated as indicated with anti-GFP RNAi and then analysed by time-lapse video microscopy. Arrows indicate the persistence of mCherry-PSF-1 on condensed chromosomes after depletion of GFP-SLD-2. PB = Polar body, located below the condensed chromosomes. (**E**) Analogous experiment with worms expressing SLD-2-GFP and mCherry-PSF-1. The arrows indicate persistence of mCherry-PSF-1 on condensed chromosomes after depletion of GFP-SLD-2. All scalebars correspond to 5µm.

We previously showed that *mcm-10Δ* is synthetic lethal in combination with incomplete depletion by RNA interference (RNAi) of the essential DNA replication initiation factors SLD-2, DNSN-1, or MUS-101 that is the worm orthologue of yeast Dpb11 and mammalian TOPBP1 (Xia *et al*., 2023). Therefore, SLD-2, DNSN-1 and MUS-101 are all candidates for factors that contribute to CMG helicase activation in the absence of MCM-10. However, DNSN-1 and MUS-101 are essential for assembly of the CMG helicase (Xia *et al*., 2023), making it harder to address whether they also have a subsequent role in helicase activation. Until now, SLD-2 was also thought to be important for CMG assembly in *C. elegans* (Gaggioli *et al*, 2014; Xia *et al*., 2023), analogous to the reported role of yeast Sld2 (Muramatsu *et al*., 2010; Yeeles *et al*., 2015). However, previous studies of worm SLD-2 did not monitor CMG assembly during S-phase and instead relied on indirect proxies such as the level of chromatin-bound CMG during mitosis of the first embryonic cell cycle, following inactivation of the CDC-48_NPL-4_UFD-1 unfoldase that normally disassembles CMG during DNA replication termination (Sonneville *et al*., 2017).

To study the role of SLD-2 in assembly of the CMG helicase during the first embryonic cell cycle, we treated *sld-2-GFP* worms with anti-GFP RNAi. In contrast to *sld-2* RNAi that depleted SLD-2-GFP incompletely (Xia *et al*., 2023), anti-GFP RNAi led to very efficient depletion of SLD-2-GFP (Figure 1D-E) and caused embryonic lethality (Figure EV1C), mirroring the lethal effects of deleting the *sld-2* gene (Xia *et al*., 2023). When embryos were treated with a short pulse of EdU at the start of S-phase, DNA synthesis was reduced about four-fold in the absence of SLD-2 compared to the control (Figure 1B-C). In addition, chromosome decondensation was delayed when embryos lacking SLD-2 entered S-phase (Figure 1D), and GINS was recruited to the condensed chromosomes during this period (Figure 1E), mirroring the situation in *mcm-10Δ* worms (Figure 1A). In contrast, GINS was not detected on chromatin when assembly of the CMG helicase was blocked and chromosome decondensation delayed by depletion of DNSN-1 (Xia *et al*., 2023). These data suggest that SLD-2 and MCM-10 are dispensable for CMG helicase assembly during the initiation of DNA replication in *C.elegans*. Instead, both MCM-10 and SLD-2 are important, though not individually essential, for activation of the CMG helicase.

Although DNA replication is delayed during the first embryonic cell cycle in the absence of SLD-2 (Figure 1B-C), chromosome condensation and segregation in the subsequent mitosis proceed normally (Figure EV2A, GFP RNAi), indicating that chromosome replication is largely completed before the initiation of chromosome segregation. In the second cell cycle of embryos lacking SLD-2, mitosis also proceeded normally in the ‘P1 cell’ (Figure EV1A-B; Figure EV2B, GFP RNAi), which is the precursor of the germline and has a longer cell cycle than the ‘AB cell’ that gives rise to the majority of the somatic cells. However, chromosome condensation was defective and chromosome segregation failed in the AB cell in all embryos lacking SLD-2 (Figure EV2C, GFP RNAi, n = 5), likely due to the shorter cell cycle of the AB cell that does not give enough time to complete replication, when CMG activation is delayed in the absence of SLD-2. These data explain why efficient depletion of SLD-2 leads to a complete loss of embryonic viability (Figure EV1C), and deletion of the *sld-2* gene is lethal in *C. elegans* (Xia *et al*., 2023). In contrast, mitosis proceeds normally in the absence of MCM-10 (Figure EV2), indicating that the initiation of DNA replication is delayed more severely in the absence of SLD-2, compared to worms lacking MCM-10.

Since CMG activation is impaired but not completely blocked upon depletion of SLD-2, some origins still fire and produce forks that slowly replicate the remainder of the genome. In this way, inactive CMG complexes at unfired origins are likely to be displaced during passive replication, by forks from neighbouring origins, thereby explaining the low level of chromatin-bound CMG during mitosis that was previously reported when depletion of SLD-2 was combined with inactivation of the CDC-48_NPL-4_UFD-1 unfoldase (Xia *et al*., 2023).

### MCM-10 and SLD-2 associate with the CMG helicase on chromatin during DNA replication initiation

Consistent with MCM-10 playing a role during CMG helicase activation *in vivo*, GFP-MCM-10 associated with chromatin during early S-phase (Figure 2A, Control), dependent upon CDC-45 (Figure 2A, *cdc-45* RNAi). In contrast, the CMG assembly factor DNSN-1 remained on condensed chromosomes in the absence of CDC-45 (Figure 2B), as reported previously (Xia *et al*., 2023), since DNSN-1 is part of a ‘pre-initiation complex’ that represents an intermediate en route to CMG assembly. However, when CMG activation and chromosome decondensation were delayed by depletion of SLD-2, GFP-MCM-10 remained on the condensed chromosomes (Figure 2A, *sld-2* RNAi). This indicates that MCM-10 is recruited to nascent CMG helicase complexes during the initiation of DNA replication in *C. elegans*, independently of SLD-2. Consistent with this view, MCM-10 binds directly to recombinant *C. elegans* CMG helicase complexes *in vitro* (Figure 2C).

**Figure 2.**
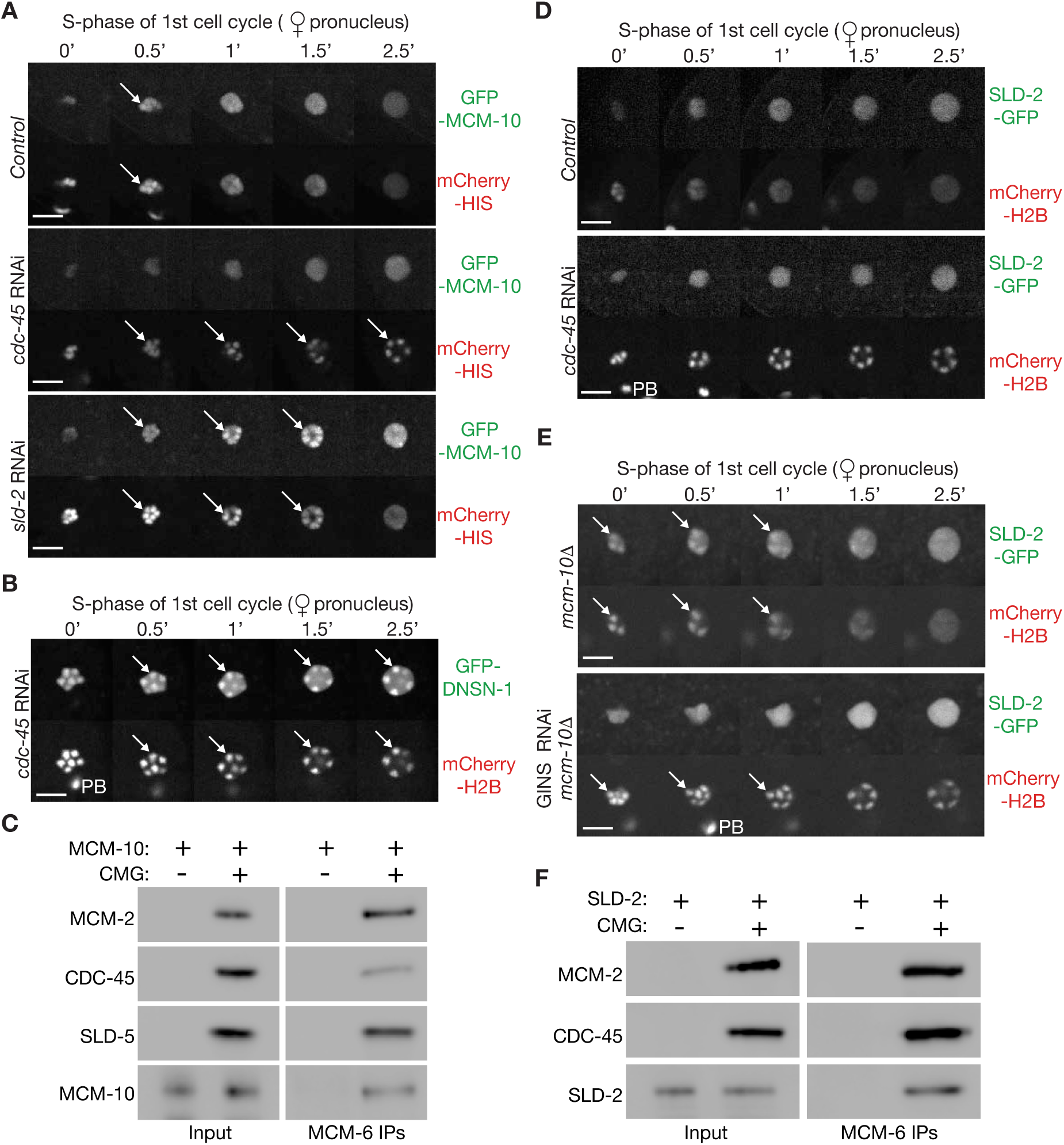
MCM-10 and SLD-2 are recruited to the CMG helicase during early S-phase in the *C. elegans* embryo. (**A**) Worms expressing GFP-MCM-10 and mCherry-HistoneH2B were treated with the indicated RNAi and then analysed by time-lapse video microscopy. Arrows indicate GFP-MCM-10 and mCherry-HistoneH2B on condensed chromosomes. (**B**) Worms expressing GFP-DNSN-1 and mCherry-HistoneH2B were treated with *cdc-45* RNAi and then analysed as above. Arrows indicate GFP-DNSN-1 and mCherry-HistoneH2B on condensed chromosomes. PB = Polar body, located below the condensed chromosomes. (**C**) Recombinant *C. elegans* MCM-10 and CMG (see Methods) were mixed *in vitro* as indicated, before isolation of CMG via immunoprecipitation of the MCM-6 subunit. The indicated proteins were detected by immunoblotting. (**D**) Worms expressing SLD-2-GFP and mCherry-HistoneH2B were treated as shown and then analysed as above. (**E**) *mcm-10Δ* worms expressing SLD-2-GFP and mCherry-HistoneH2B were treated as shown with RNAi to the four subunits of GINS, before analysis by time-lapse video microscopy. Arrows indicate SLD-2-GFP and mCherry-HistoneH2B on condensed chromosomes. (**F**) Recombinant *C. elegans* SLD-2 and CMG (see Methods) were processed as in (C). All scalebars correspond to 5µm.

In wild type embryos, SLD-2 is not detected on chromosomes during early S-phase (Figure 2D, Control), suggesting that its association with CMG is very transient. Furthermore, and analogous to the behaviour of MCM-10 (Figure 2A, *cdc-45* RNAi), SLD-2 does not accumulate on chromosomes when CMG assembly is blocked by depletion of CDC-45 (Figure 2D, *cdc-45* RNAi). However, SLD-2-GFP was detected on condensed chromosomes when CMG activation was delayed in the absence of MCM-10 (Figure 2E, *mcm-10Δ*; also see Figure 3A & 3C, *mcm-10Δ*). Moreover, the chromatin association of SLD-2 under such conditions was blocked by depletion of GINS (Figure 2E, GINS RNAi *mcm-10Δ*). These data indicate that SLD-2 associates transiently with newly assembled CMG helicase complexes during the initiation of DNA replication, independently of MCM-10. Correspondingly, SLD-2 can bind directly to the CMG helicase *in vitro* (Figure 2F).

**Figure 3.**
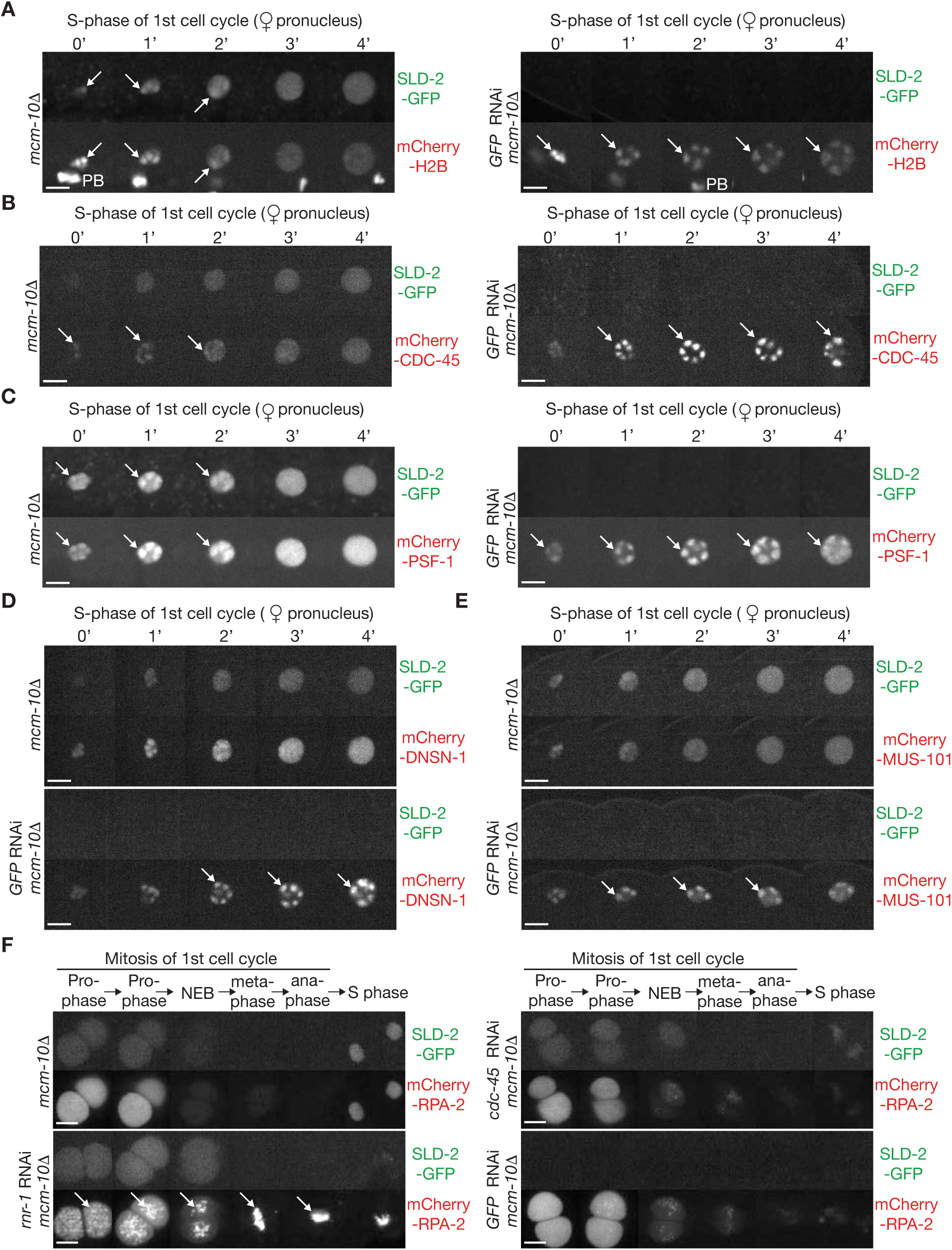
CMG helicase activation is blocked in the combined absence of *C. elegans* MCM-10 and SLD-2. (**A**) Worms of the genotype *mcm-10Δ sld-2-GFP mCherry-histoneH2B* were treated as indicated with anti-GFP RNAi, then processed as for Figure 1D. Arrows indicate SLD-2-GFP and mCherry-HistoneH2B on condensed chromosomes. PB = Polar body, located below the condensed chromosomes. (**B**) Similar experiment with *mcm-10Δ sld-2-GFP mCherry-cdc-45*. Arrows indicate the association of mCherry-CDC-45 with condensed chromosomes in worms lacking both SLD-2 and MCM-10. Note that SLD-2-GFP does not accumulate on condensed chromosomes in *mcm-10Δ* worms expressing *mCherry-CDC-45* (top panel), likely due to partial impairment of CDC-45 function upon fusion to the mCherry tag. (**C**-**E**) Analogous experiments with *mcm-10Δ sld-2-GFP mCherry-psf-1* (C), *mcm-10Δ sld-2-GFP mCherry-dnsn-1* (D) or *mcm-10Δ sld-2-GFP mCherry-mus-101* (E). Arrows show the association of mCherry-PSF-1 (C), mCherry-DNSN-1 (D) and mCherry-MUS-101 (E) with condensed chromosomes in embryos lacking both SLD-2 and MCM-10. (**F**) *mcm-10Δ* worms expressing SLD-2-GFP and mCherry-RPA-2 were treated with RNAi as indicated, before analysis by time-lapse video microscopy of progression through mitosis in the first embryonic cell cycle. Arrows indicate SLD-2-GFP and mCherry-HistoneH2B on condensed chromosomes. All scalebars correspond to 5µm.

### MCM-10 and SLD-2 are jointly required for CMG helicase activation in *C. elegans*

Depletion of SLD-2 in *mcm-10Δ* worms strongly inhibited DNA replication in the first embryonic cell cycle, compared to embryos that individually lacked either MCM-10 or SLD-2 (Figure 1B-C). Moreover, chromosome decondensation was delayed for longer in embryos lacking both MCM-10 and SLD-2, compared to the effect of removing just one of the two proteins (Figure 3A, Figure 1D). Both CDC-45 (Figure 3B) and GINS (Figure 3C) accumulated on condensed chromosomes in the absence of MCM-10 and SLD-2. Similarly, the initiation factors DNSN-1 (Figure 3D) and MUS-101 (Figure 3E) persisted on condensed chromosomes in embryos lacking MCM-10 and SLD-2. Collectively, these findings indicated that the CMG helicase was assembled in the absence of MCM-10 and SLD-2 but remained in a ‘pre-activation complex’ with DNSN-1 and MUS-101, rather than giving rise to active replisomes. Consistent with a defect in DNA unwinding, the single-strand DNA-binding protein RPA (Replication Protein A) did not accumulate on chromatin in embryos lacking MCM-10 and SLD-2 (Figure 3F, GFP RNAi + *mcm-10Δ*). This situation resembled the effect of blocking CMG assembly by depleting CDC-45 (Figure 3F, *cdc-45* RNAi + *mcm-10Δ*), whereas RPA accumulated on chromatin when DNA synthesis was inhibited after CMG helicase activation, by depletion of the ribonucleotide reductase subunit RNR-1 (Figure 3F, *rnr-1* RNAi + *mcm-10Δ*). Collectively, these findings indicated that MCM-10 and SLD-2 are jointly required for CMG helicase activation, which is profoundly defective in their absence. Correspondingly, chromosome condensation during mitotic prophase of the first embryonic cell cycle was impaired when SLD-2 was depleted in *mcm-10Δ* worms, and chromosome segregation failed during anaphase, producing chromatin bridges in all embryos (Figure EV2A, GFP RNAi + *mcm-10Δ*, n = 5).

### MCM-10, SLD-2 and DNSN-1 stimulate CMG helicase activity *in vitro*

Given the evidence that MCM-10 and SLD-2 are important for CMG helicase activation *in vivo*, we tested whether they can directly stimulate CMG helicase activity *in vitro*, as reported previously for budding yeast Mcm10 (Langston *et al*, 2017; Langston & O’Donnell, 2019; Looke *et al*, 2017; Wasserman *et al*, 2019). MCM-10 and SLD-2 were expressed in recombinant form in budding yeast cells and then purified from cell extracts, as were MUS-101 and DNSN-1 that also promote the initiation of DNA replication in *C. elegans* (Figure 4A). We also expressed and purified a set of proteins that bind to the CMG helicase in the *C. elegans* replisome (Sonneville *et al*., 2017; Xia *et al*., 2021), such as the TIM-1_TIPN-1 complex (TIMELESS-TIPIN), CLSP-1 (CLASPIN), CTF-4 and the CTF-18_RFC clamp loader complex. Finally, we purified the single-strand DNA binding factor known as Replication Protein A (RPA), which is a two-protein complex in *C. elegans* rather than three subunits in many other eukaryotic species. Subsequently, we assayed the ability of each factor to stimulate the ability of the *C. elegans* CMG helicase to unwind a model DNA replication substrate, in which one of the two strands was radioactively labelled (Figure 4B).

**Figure 4.**
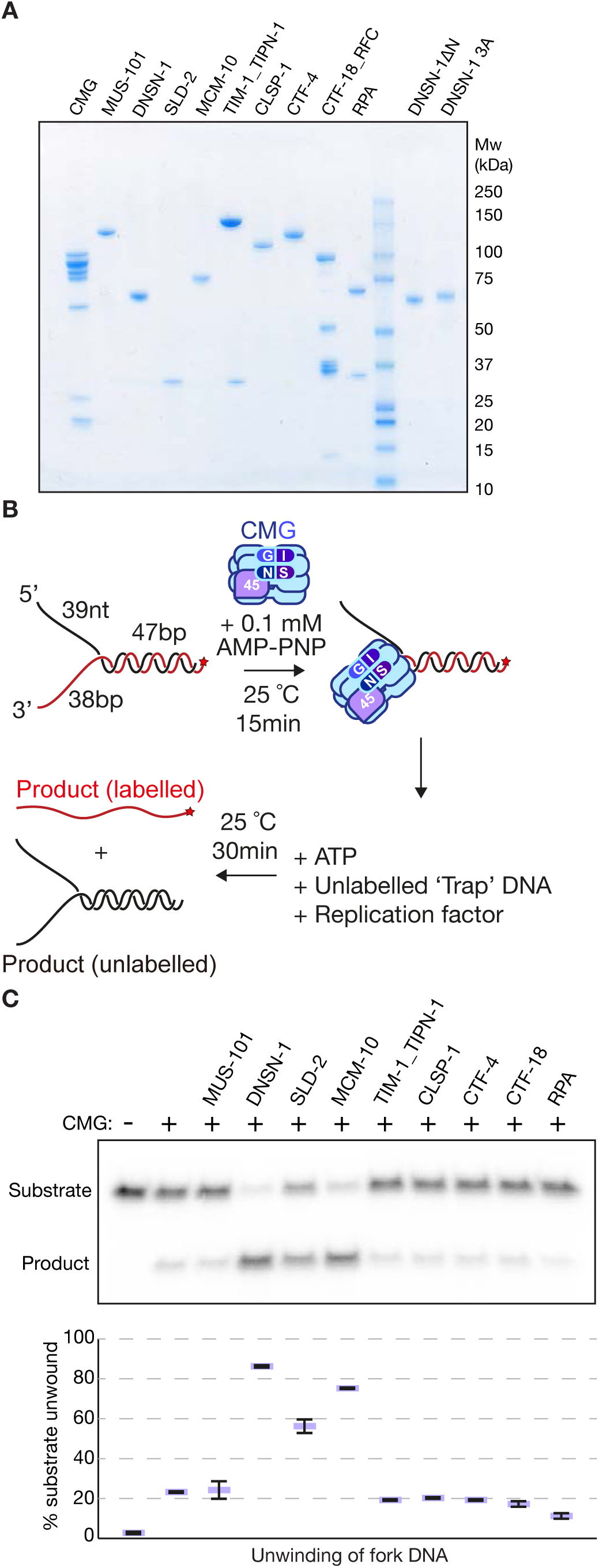
MCM-10, SLD-2 and DNSN-1 stimulate activity of the CMG helicase in vitro. (**A**) The indicated factors were expressed and purified in recombinant form as described in Methods. The purified proteins were resolved by SDS-PAGE and stained with colloidal Coomassie blue. (**B**) Reaction scheme (radio-labelling of 5’-end of DNA strand is indicated by a red star). (**C**) Helicase assays were performed as in (B) with the factors shown in (A). Reaction products were separated in a polyacrylamide gel and detected by autoradiography. The percentage of unwound substrate DNA was quantified and shown in the graph at the bottom, indicating mean values and standard deviation from three independent experiments.

In contrast to the other tested factors, MCM-10, SLD-2 and DNSN-1 all stimulated CMG helicase activity *in vitro* (Figure 4C and Figure EV3). All three proteins share the ability to bind directly to the CMG helicase (Figure 2C+F and (Xia *et al*., 2023)) and to DNA (Figure EV4). Whereas the interaction of metazoan MCM-10 with CMG is poorly characterised and the binding of SLD-2 to CMG has not been studied structurally, the association of DNSN-1 with CMG has been analysed by cryo-electron microscopy, revealing the importance of an amino-terminal helix in DNSN-1 that binds to the SLD-5 subunit of GINS, and three conserved DNSN-1 residues that bind to MCM-3 (Xia *et al*., 2023). Removal of the amino-terminal helix prevents DNSN-1 from binding to CMG (Figure EV5A), as seen previously (Xia *et al*., 2023), and impaired the ability of DNSN-1 to stimulate CMG helicase activity *in vitro* (Figure EV5B). Furthermore, mutation of the MCM-3-binding site led to partial displacement of DNSN-1 from CMG (Figure EV5A), and partially reduced the ability of DNSN-1 to stimulate CMG helicase activity (Figure EV5B). These findings indicate that *C. elegans* MCM-10, SLD-2 and DNSN-1 all stimulate CMG helicase activity *in vitro*, most likely via direct binding to CMG and to DNA.

### Inactivation of MCM10 and RECQL4 causes synthetic lethality in mouse ES cells

Previous studies showed that transient depletion of MCM10 in human cancer cell lines delayed cell cycle progression, probably due to defects in DNA replication (Chattopadhyay & Bielinsky, 2007). Moreover, deletion of the mouse *Mcm10* gene causes lethality around day 8.5 of embryonic development (Lim *et al*, 2011). These findings suggested that MCM10 is essential for DNA replication in mammalian cells.

In contrast to yeasts, the orthologue of Sld2 in many eukaryotic species contains a DNA helicase of the RecQ family, after the region of Sld2 homology, making a protein that in vertebrates is called RECQL4 (Gaggioli *et al*., 2014). Studies of RECQL4 in *Xenopus laevis* indicated that it is essential for the initiation of DNA replication, dependent upon the amino-terminal region that is homologous to yeast Sld2 (Matsuno *et al*, 2006; Sangrithi *et al*, 2005). However, unlike Sld2 in *Saccharomyces cerevisiae*, *Xenopus* RECQL4 is dispensable for CMG assembly and instead is required for helicase activation (Matsuno *et al*., 2006; Sangrithi *et al*., 2005). Nevertheless, recent studies showed that human cells can survive in the absence of detectable expression of RECQL4 (Padayachy *et al*, 2024). Small deletions or insertions in exon 1 of RECQL4 were introduced by CRISPR-Cas9 genome editing in human U2OS cells, producing viable cells in which expression of the RECQL4 protein was not detected, although it remains possible that truncated forms of the protein were still expressed at a low level and are important to preserve viability (Padayachy *et al*., 2024). Indeed, a very recent study of mouse myeloid cells showed that cell viability could be maintained by expression of an amino-terminal fragment of RECQL4, containing the region of homology to yeast and *C. elegans* Sld2 / SLD-2, whereas complete loss of RECQL4 was lethal (Buco *et al*, 2025).

To explore whether the joint role of *C. elegans* MCM-10 and SLD-2 in the activation of the CMG helicase is conserved in mammalian cells, we inactivated the *Mcm10* and *Recql4* genes in mouse embryonic stem cells (mouse ES cells), which spend most of the cell cycle in S-phase and provide a model system for studying the regulation of the mammalian CMG helicase (Evrin *et al*., 2023; Villa *et al*, 2021). Firstly, we used CRISPR and the Cas9-D10A nickase enzyme, together with pairs of guide RNAs that target exon 3 of *Mcm10* (Figure EV6A) or exon 1 of *Recql4* (Figure EV6B), to make small deletions at the 5’ end of the coding sequence. In this way, we isolated viable clones that lacked detectable expression of MCM10 or RECQL4 (Figure EV6C-F). *Mcm10Δ* clones grew at a similar rate to control cells, whereas *Recql4Δ* cells grew more slowly (19.9 +/− 1.8 hours for *Recql4Δ*, compared to 13 +/−0.5 hours for control).

Subsequently, we used CRISPR and wild type Cas9 to cut both ends of the *Mcm10* coding sequence and delete the entire 22.6 kb gene (Figure EV7A-C). This produced viable clones that lacked MCM10 protein (Figure EV7D) and grew slightly slower than control cells (15 +/− 0.7 hours). These data show that mouse ES cells are able to grow in the complete absence of MCM10, indicating that mammalian cells must have at least one MCM10-independent mechanism for activating the CMG helicase. We used a similar approach to try and delete the entire 7 kb *Recql4* gene (Figure EV8), but only isolated clones that were heterozygous for complete deletion of *Recql4*, suggesting that low-level expression of truncated fragments of RECQL4 is important to preserve cell viability in mouse ES cells.

Since mouse ES cells are able to grow in the absence of MCM10 and with an extremely low level of RECQL4 (below the detection limit in immunoblots), we tested the effect of using CRISPR-Cas9 to make deletions in both genes simultaneously, using the approach in Figure EV6. Whereas *Mcm10Δ* had little impact on cell growth in control cells, the combination of *Mcm10Δ* with *Recql4Δ* inhibited colony formation (Figure 5A). Moreover, *Recql4Δ* had a modest impact on colony formation in control cells but strongly impaired the growth of *Mcm10Δ* cells (Figure 5B). These findings indicated that simultaneous loss of MCM10 and RECQL causes synthetic lethality in mouse ES cells.

**Figure 5.**
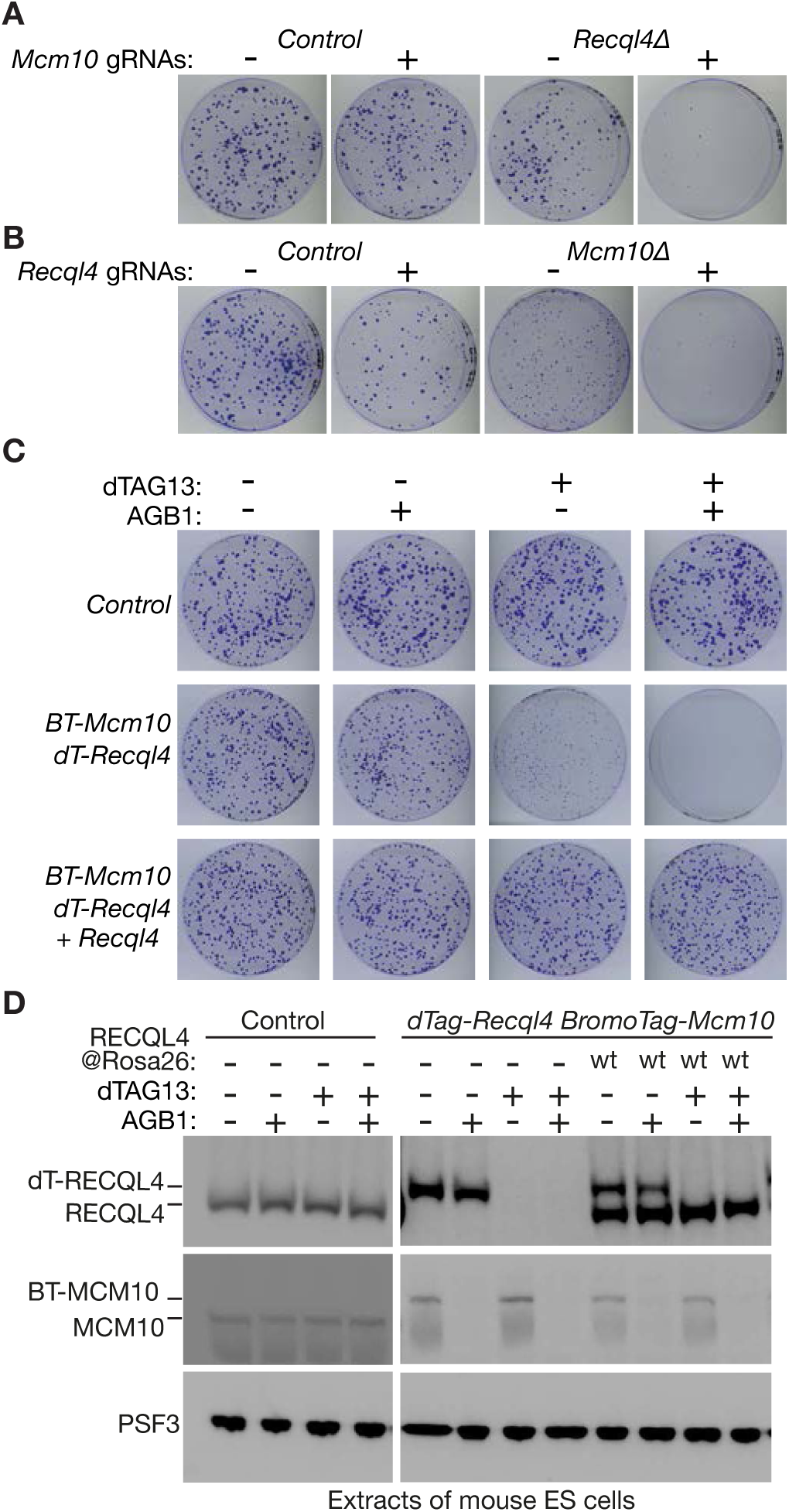
Combined depletion of MCM10 and RECQL4 causes synthetic lethality in mouse embryonic stem cells. (**A**) Control mouse ES cells and *Recql4Δ* cells (Figure EV6) were transfected as indicated with plasmids expressing the Cas9-D10A nickase and guide RNAs (gRNAs) specific to exon 3 of the murine *Mcm10* gene (Figure EV6A). Cell proliferation was then assayed as described in Methods. (**B**) Similar experiment to (A) but with control mouse ES cells and *Mcm10Δ* cells (Figure EV6) transfected with plasmids expressing the Cas9-D10A nickase and guide RNAs (gRNAs) specific to exon 1 of the murine *Recql4* gene (Figure EV6B). (**C**) Mouse ES cells of the indicated genotypes (BT = BromoTag; dT = dTAG; ‘+ Recql4’ = expression of wild type *Recql4* from *Rosa26* locus) were treated as shown with the PROTACs AGB1 or dTAG13, before monitoring cell proliferation as described in Methods. (**D**) Expression of the indicated proteins was monitored by immunoblotting in the experiment shown in (C).

To study the phenotype of mammalian cells lacking MCM10 and RECQL4 in more depth, we generated a conditional depletion system. CRISPR-Cas9 was used to make mouse ES cells in which MCM10 was fused to BromoTag that is recognised by the ubiquitin ligase CUL2^VHL^ in the presence of the Proteolysis Targeting Chimera (PROTAC) known as AGB1 (Bond *et al*, 2021). In addition, RECQL4 was fused to dTAG (Nabet *et al*, 2018) that is recognised by the ubiquitin ligase CUL4^CRBN^ in the presence of a PROTAC called dTAG13 (Figure EV9). The growth of control cells was not affected by exposure to either or both of the two PROTACs (Figure 5C, Control). Moreover, *BromoTag-Mcm10 dTAG-Recql4* cells grew well when BromoTag-MCM10 was depleted by addition of AGB1, and more slowly after depletion of dTAG-RECQL4 by addition of dTAG13 (Figure 5C-D, *BromoTag-Mcm10 dTAG-Recql4*), consistent with the knockout data described above. However, simultaneous depletion of both BromoTag-MCM10 and dTAG-RECQL4 was lethal (Figure 5C-D, *BromoTag-Mcm10 dTAG-Recql4*, AGB1+dTAG13). Furthermore, this effect could be rescued by expression of wild type RECQL4 from the *Rosa26* locus (Figure 5C-D, *BromoTag-Mcm10 dTAG-Recql4* + *Recql4*, AGB1+dTAG13). These findings confirm that simultaneous loss of MCM10 and RECQL4 prevents the proliferation of mouse ES cells.

### Absence of MCM10 and RECQL4 blocks DNA replication after assembly of the CMG helicase

To examine the impact of depleting MCM10 and RECQL4 on cell cycle progression, we synchronised *BromoTag-Mcm10 dTAG-Recql4* cells in S-phase, released them into the subsequent mitosis, and then held them in mitosis for a further period, in the presence or absence of the PROTACs AGB1 and dTAG13, before releasing cells from mitotic arrest (Figure 6A). Depletion of MCM10 and RECQL4 (Figure 6B) had no impact on progression through mitosis or cell division, but inhibited DNA replication in the subsequent cell cycle (Figure 6A). By immunoprecipitation of a GFP-tagged form of the GINS subunit PSF1, we observed that cells expressing MCM10 and RECQL4 assembled the CMG helicase complex upon entry into S-phase (Figure 6C, -PROTACs). Importantly, cells lacking MCM10 and RECQL4 also assembled the CMG helicase upon entry into S-phase (Figure 6C, +PROTACs). These findings indicate that MCM10 and RECQL4 both contribute to an essential step during chromosome replication in the mammalian cell cycle, downstream of assembly of the CMG helicase.

**Figure 6.**
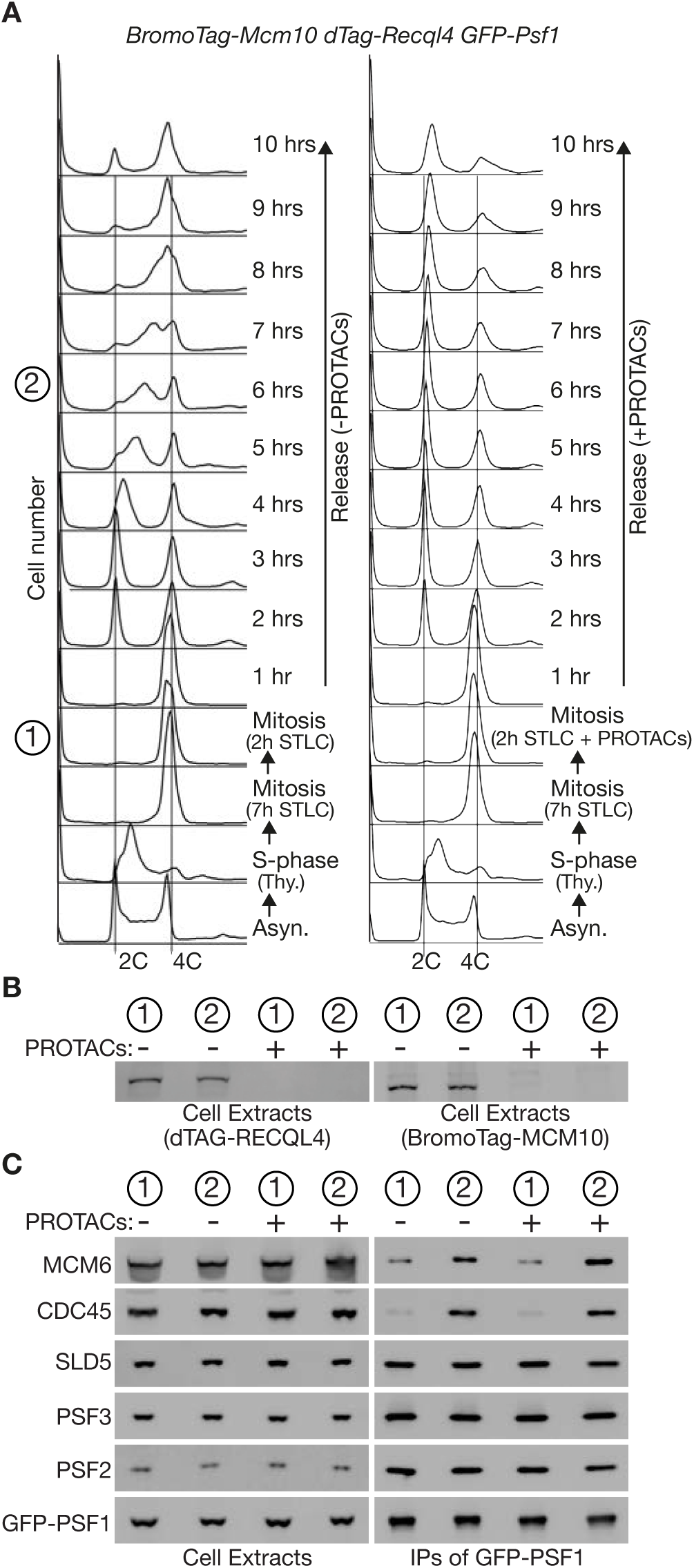
MCM10 and RECQL4 are jointly required for DNA replication after CMG helicase assembly in mouse ES cells. (**A**) Mouse ES cells of the indicated genotype were synchronised firstly in S-phase by addition of thymidine, and then in mitosis in the presence of the kinesin inhibitor STLC, as described in Methods. Half of the culture was then exposed for two hours to the PROTACs AGB1 and dTAG13, to induce degradation of BromoTag-MCM10 and dTAG-RECQL4, before cells were released from mitosis and allowed to progress into the subsequent cell cycle. DNA content was monitored throughout the experiment by flow cytometry, and additional samples were taken at the end of the mitotic arrest (‘1’) and six hours after release from mitosis (‘2’), in order to monitor protein expression and assembly of the CMG helicase. (**B**) The level of BromoTag-MCM10 (left) and dTAG-RECQL4 (right) was monitored by immunoblotting at the indicated stages of the experiment in (A). (**C**) Native protein extracts were generated at the indicated times from the experiment in (A) and used to isolate the GINS sub-complex (SLD5-PSF1-PSF2-PSF3) of the CMG helicase, by immunoprecipitation of GFP-tagged PSF1. The indicated proteins were monitored by immunoblotting.

### CMG helicase activation is defective in mammalian cells lacking MCM10 and RECQL4

To examine in more detail the impact of entering S-phase in the absence of MCM10 and RECQL4, we used high-content screening microscopy to analyse the behaviour of thousands of individual cells, and compared the impact of degrading MCM10 and RECQL4 with degradation of the essential CMG assembly factor DONSON. Consistent with previous findings (Evrin *et al*., 2023), degradation of DONSON in mouse ES cells (Figure 7A) impaired DNA synthesis (Figure 7B), did not prevent association of the MCM2-7 proteins with chromatin during G1-phase (Figure 7C), but blocked the recruitment of CDC45 (Figure 7D) and GINS (Figure 7E) to chromatin during S-phase. Depletion of both MCM10 and RECQL4 (Figure 7F, lane 8) inhibited DNA synthesis to a similar degree to loss of DONSON (Figure 7G) and also did not affect loading of MCM2-7 proteins on chromatin during G1-phase (Figure 7H). However, chromatin association of CDC45 (Figure 7I) and GINS (Figure 7J) was enhanced rather than impaired, in S-phase cells that lacked MCM10 and RECQL4. This novel phenotype was not seen when the progression of DNA replication forks was blocked by treating cells with hydroxyurea that inhibits ribonucleotide reductase and impairs the synthesis of dNTPs (Figure 8A-C), suggesting that enhanced loading of CDC45 and GINS does not result from a defect in the elongation stage of chromosome replication. Instead, these data suggested that the CMG helicase assembles during DNA replication initiation in the absence of MCM10 and RECQL4, but remains inactive, leading to CMG assembly at progressively more origins in S-phase cells, without being balanced by the progression and termination of DNA replication forks that normally triggers disassembly of the CMG helicase in control cells (Dewar *et al*, 2017; Jenkyn-Bedford *et al*, 2021; Jones *et al*, 2024; Low *et al*, 2020; Villa *et al*., 2021).

**Figure 7.**
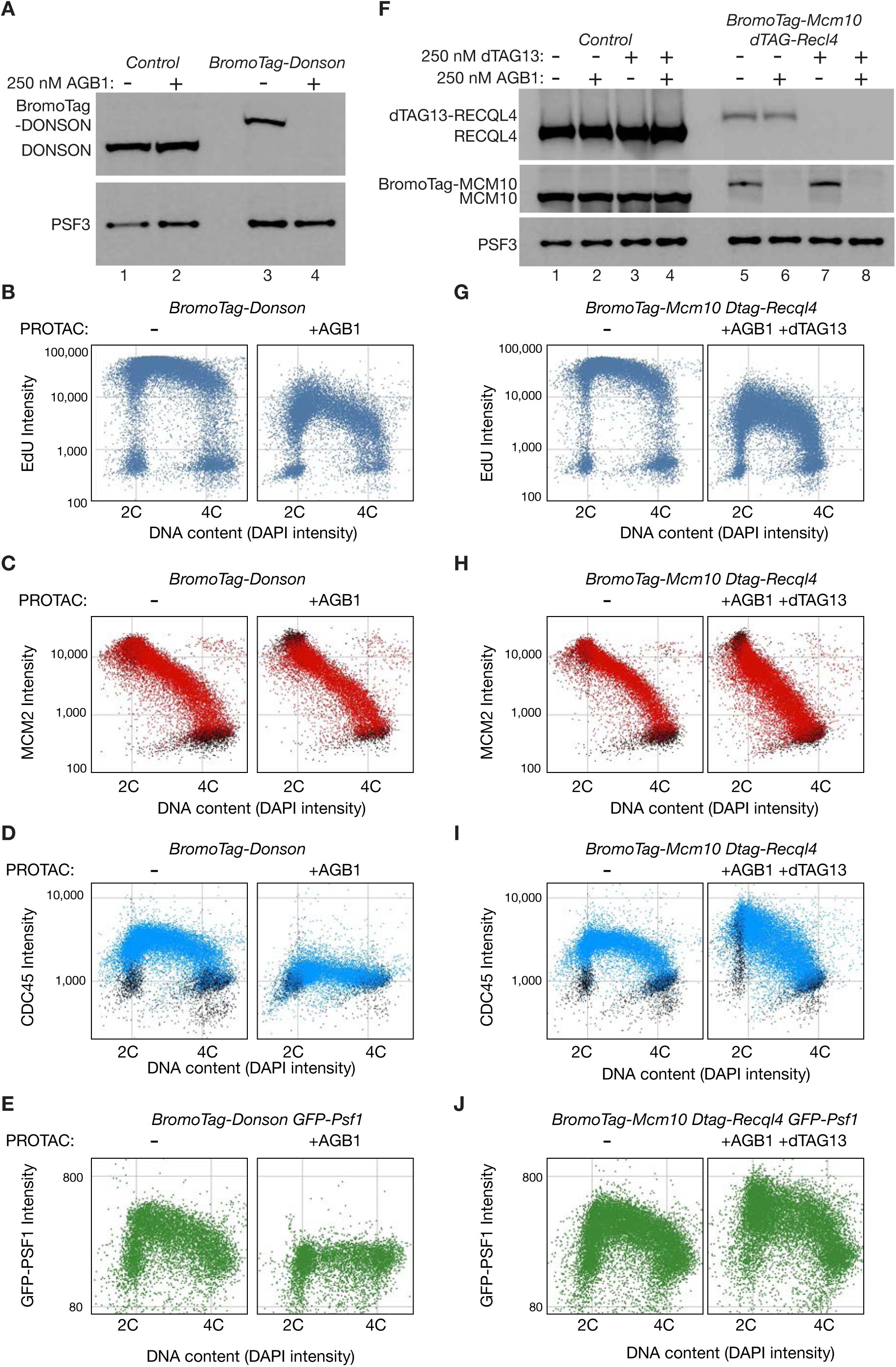
The CMG helicase accumulates on chromatin in mouse ES cells lacking MCM10 and RECQL4. (**A**) Asynchronous cultures of control and *BromoTag-Donson* mouse ES cells were incubated for three hours as indicated in the presence or absence of the PROTAC AGB1. Cell extracts were used to monitor the level of the indicated proteins by immunoblotting. (**B** - **E**) *BromoTag-Donson* mouse ES cells were incubated for eight hours in the presence or absence of AGB1, followed by treatment for 20 minutes with the nucleoside precursor EdU. Cells were then analysed by high-content screening microscopy as described in Methods, to monitor DNA synthesis (EdU intensity) and chromatin association of the indicated components of the CMG helicase. In contrast to control cells (Figure EV10), depletion of *BromoTag-Donson* impairs DNA synthesis and CMG helicase assembly, as described previously (Xia *et al*., 2023). For panels (C) and (D), EdU-negative cells (see Methods) are indicated in black. For panel (E), cells also expressed GFP-PSF1. (**F**) Control and *BromoTag-Mcm10 dTAG-Recql4* mouse ES cells were incubated for three hours as indicated in the presence or absence of the PROTACs AGB1 and dTAG13. Cell extracts were used to monitor the level of the indicated proteins by immunoblotting. (**G** - **J**) *BromoTag-Mcm10 dTAG-Recql4* mouse ES cells were incubated for eight hours in the presence or absence of AGB1 and dTAG13, followed by treatment for 20 minutes with EdU. Cells were then analysed as described above by high-content screening microscopy. For panels (H) and (I), EdU-negative cells (see Methods) are indicated in black. For panel (J), cells also expressed GFP-PSF1. Equivalent data for control cells, and for *BromoTag-Mcm10 dTAG-Recql4* cells in which just one of BromoTag-MCM10 or dTAG-RECQL4 was depleted, are shown in Figure EV10.

**Figure 8.**
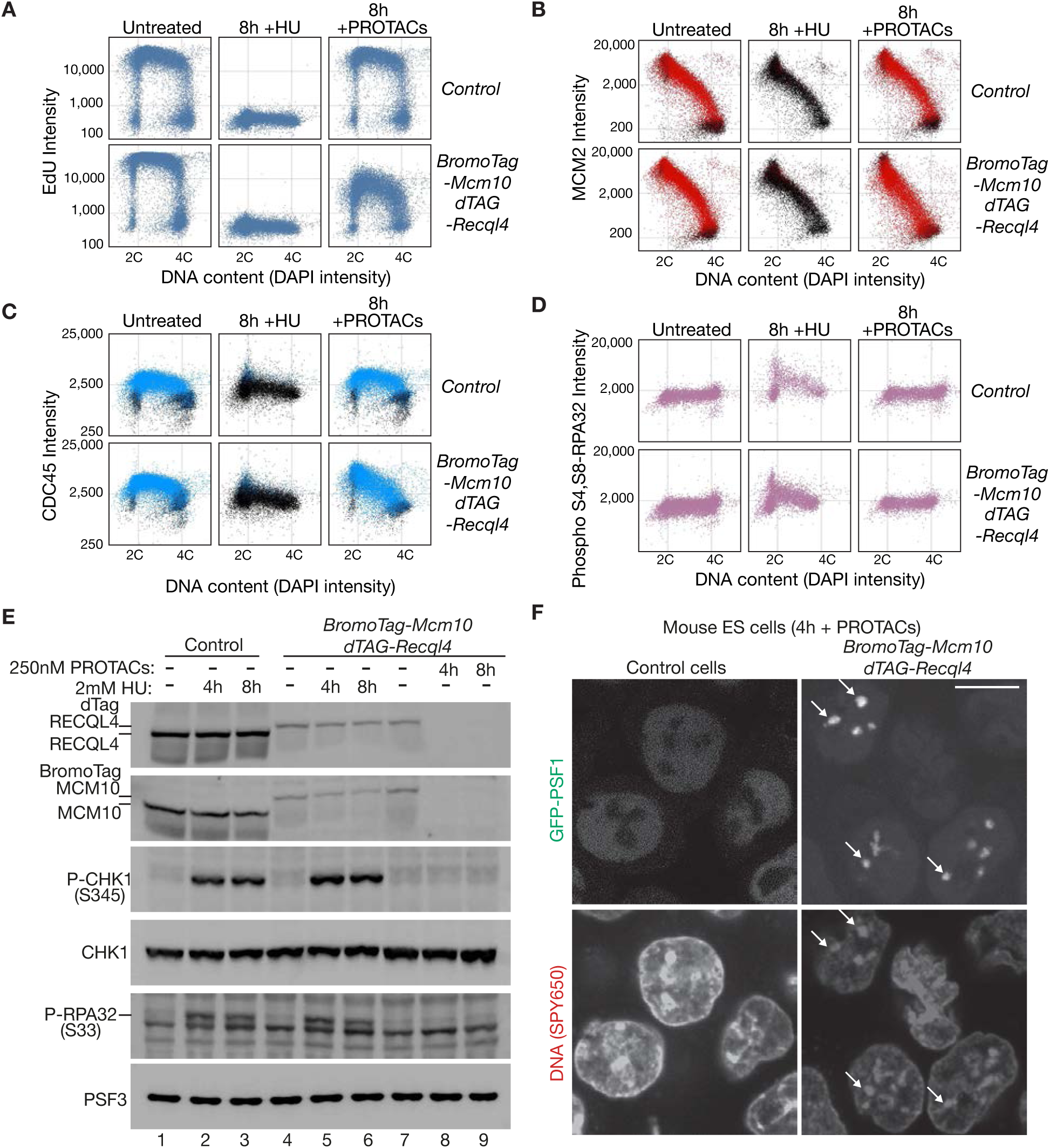
The DNA replication checkpoint is inactive in the absence of MCM10 and RECQL4, indicating a failure to activate the CMG helicase. (**A** - **C**) Control and *BromoTag-Mcm10 dTAG-Recql4* mouse ES cells were incubated for eight hours in the presence or absence of the PROTACs AGB1 and dTAG13, or in the presence of 2 mM hydroxyurea, before treatment for 20 minutes with EdU. Cells were analysed by high-content screening microscopy as described above for Figure 7. For panels (B) and (C), EdU-negative cells (see Methods) are indicated in black. (**D**) Samples for the above high-content screening experiment were also analysed by immuno-staining with phospho-specific antibodies corresponding to Serine 4 (S4) and Serine 8 (S8) of RPA32, which is phosphorylated in response to activation of the DNA replication checkpoint pathway. (**E**) Samples from the same experiment were used to prepare cell extracts that were analysed by immunoblotting for the indicated proteins. ‘P-CHK1 (S345)’ corresponds to phosphorylation of Serine 345 of CHK1, whereas ‘P-RPA32 (S33)’ corresponds to phosphorylation of Serine 33 of RPA32, both of which are mediated by the ATR kinase in response to checkpoint activation. (**F**) Mouse ES cells of the genotype *GFP-Psf1* (Control) or *BromoTag-Mcm10 dTAG-Recql4 GFP-Psf1* were incubated for four hours in the presence or absence of the PROTACs AGB1 and dTAG13. Live cells were then stained with the DNA-binding dye SPY650-DNA (see Methods). Arrows indicate the accumulation of GFP-PSF1 on patches of constitutive heterochromatin. The scale bar corresponds to 10 µm.

To test whether DNA unwinding occurs during the initiation of DNA replication, when cells lacking MCM10 and RECQL4 assemble CMG but fail to synthesise their DNA, we monitored activation of the signalling pathway known as the DNA replication checkpoint (Yates *et al*, 2025). When DNA synthesis is blocked in cells with active CMG helicase, DNA unwinding at stalled DNA replication forks leads to activation of the checkpoint. The ATR kinase is recruited to single-strand DNA that is coated by RPA at stalled replication forks, leading to activation of ATR, which then phosphorylates and activates the downstream kinase CHK1. ATR and CHK1 then inhibit proteins that are required for CMG helicase assembly (Boos *et al*, 2011; Lee *et al*, 2012), thereby blocking the initiation of DNA replication in later replicating parts of the genome. ATR also phosphorylates other substrates at replication forks such as the RPA subunit RPA32 on Serine 33 (Olson *et al*, 2006). In addition, RPA32 is phosphorylated on Serine 4 and Serine 8 downstream of ATR activation (Olson *et al*., 2006), by a related kinase known as DNAPK (Anantha *et al*, 2007).

Whereas treatment of control and *BromoTag-Mcm10 dTAG-Recql4* cells with hydroxyurea led to phosphorylation of RPA32-Ser4 and RPA32-Ser8 (Figure 8D, 8h +HU), RPA32-Ser33 (Figure 8E, lanes 2-3 and 5-6, P-RPA32-S33) and CHK1-Ser345 (Figure 8E, lanes 2-3 and, P-CHK1-S345), simultaneous depletion of BromoTag-MCM10 and dTAG-RECQL4 did not activate the checkpoint (Figure 8D, 8h + PROTACs; Figure 8E, lanes 8-9). Therefore, these data indicate that the CMG helicase assembles in the absence of MCM10 and RECQL4 but remains inactive. Failure to activate the checkpoint would allow the continuation of CMG assembly in later replicating regions, leading to accumulation of inactive CMG helicase complexes on chromatin. Consistent with this view, we observed the accumulation of GINS on patches of heterochromatin in cells lacking MCM10 and RECQL4 (Figure 8F; 45 % cells, n = 623, compared to 16 % of cells in the presence of MCM10 and RECQL4, n = 622). Collectively, the above data are consistent with MCM10 and RECQL4 being jointly required for activation of the CMG helicase during the initiation of DNA replication in mammalian cells.

## Discussion

When budding yeast cells enter S-phase, the phosphorylation of both Sld2 and Sld3 by Cyclin-Dependent Kinase (CDK) is essential for the initiation of chromosome replication (Tanaka *et al*, 2007; Zegerman & Diffley, 2007). Phosphorylated Sld2 and Sld3 are connected to each other by the phospho-receptor protein Dpb11, which uses pairs of BRCA1 C-Terminal (BRCT) domains to bind to phospho-peptides. Sld3 also binds to Cdc45 (Kamimura *et al*, 2001) and Mcm2-7 double hexamers (Deegan *et al*., 2016), and phospho-Sld2 forms a labile complex not only with Dpb11 but also with GINS and Pol ε (Muramatsu *et al*., 2010). Therefore, phosphorylation of Sld2 and Sld3 drives a transient network of protein-protein interactions at origins of replication, recruiting Cdc45 and GINS to Mcm2-7 double hexamers and triggering CMG helicase assembly.

The mechanism by which Sld3 recruits Cdc45 to Mcm2-7 double hexamers is likely to be conserved in diverse eukaryotes, involving orthologues of yeast Sld3 known as TRESLIN or TICRR (Kumagai *et al*, 2010; Sansam *et al*, 2010). The recruitment of GINS has diverged to a greater degree during evolution, with DONSON playing a central role in animal cells (Hashimoto *et al*., 2023; Lim *et al*., 2023; Xia *et al*., 2023), whereas Sld2 and Pol ε are important for CMG assembly in budding yeast (Muramatsu *et al*., 2010; Yeeles *et al*., 2015) but are dispensable for helicase assembly in animal cells.

The mechanism of CMG helicase activation during DNA replication initiation is still one of the least characterised steps in eukaryotic chromosome replication. Mcm10 is essential for helicase activation in budding yeast (Douglas *et al*., 2018; van Deursen *et al*., 2012; Watase *et al*., 2012) and binds both to CMG and to DNA (Gambus *et al*., 2006; Henrikus *et al*, 2024; Langston *et al*., 2017; Looke *et al*., 2017), suggesting that Mcm10 promotes helicase activation by interacting with double CMG complexes and facilitating the irreversible extrusion of one of the two DNA strands from within the central cavity of each MCM2-7 hexamer. Our data indicate that helicase activation is delayed in the absence of *C. elegans* MCM-10 (Figure 1A), but subsequently occurs since SLD-2 also promotes helicase activation during the initiation of DNA replication (Figure 1D). It will be interesting in future studies to determine whether yeast Sld2 contributes to helicase activation in addition to Mcm10, which would indicate a conserved mechanism for CMG helicase activation in diverse eukaryotic species.

It is possible that DNSN-1 also contributes to helicase activation in *C. elegans*, in addition to SLD-2 and MCM-10. Partial depletion of either DNSN-1 or SLD-2 is lethal in worms lacking MCM-10 (Xia *et al*., 2023), and our data indicate that DNSN-1 (and its partner MUS-101) associates with inactive CMG helicase complexes on chromatin in embryos lacking MCM-10 and SLD-2 (Figure 3D-E). Moreover, we show that MCM-10, SLD-2 and DNSN-1 all stimulate CMG helicase activity *in vitro* (Figure 4 and Figure EV3). All three proteins bind to CMG and DNA (Figure EV4), suggesting a common mechanism of CMG activation, whereby activator proteins bind to CMG and help to capture the DNA strand that is extruded from the central cavity of MCM-2-7. Further structural studies, together with the reconstitution of metazoan helicase activation from MCM2-7 double hexamers, will be required to investigate the mechanism in greater detail.

We show that mouse ES cells can proliferate in the absence of MCM10 (Figure EV7), mirroring the situation in *C. elegans* (Xia *et al*., 2023), and indicating the existence of MCM10-independent routes to CMG helicase activation. Mouse cells that completely lack MCM10 grow more slowly than control cells, and a delay in helicase activation likely explains why MCM10 becomes essential in some mammalian cell cycles, for example during mouse development (Lim *et al*., 2011), or in human cancer cell lines (Chattopadhyay & Bielinsky, 2007). In human patients, hypomorphic mutations in MCM10 are associated with a condition called Natural Killer Cell Deficiency, or alternatively with restrictive cardiomyopathy associated with hypoplasia of the spleen and thymus (Baxley *et al*, 2021; Mace *et al*, 2020; Schmit *et al*, 2024).

Our data indicate that RECQL4 plays a partially redundant role during CMG activation with MCM10 in mammalian cells (Figures 5-8), analogous to the situation in *C. elegans*. Moreover, our findings suggest that inactivation of RECQL4 produces a greater defect in CMG helicase activation in mouse ES cells than inactivation of MCM10 (Figure EV10C-D). This also resembles the situation in *C. elegans*, for which *mcm-10Δ* is viable (Xia *et al*., 2023), whereas absence of SLD-2 delays CMG activation and leads to the failure of nuclear division in the second cell cycle (Figure EV2C).

Consistent with previous findings in human cells (Padayachy *et al*., 2024), we were able to generate viable cells with small deletions in the mouse *Recql4* gene, which removed detectable expression of RECQL4 protein (Figure 5A-B). However, complete deletion of the *Recql4* gene was only obtained in a heterozygous form in mouse ES cells (Figure EV8). Recent work indicated that RECQL4 is essential in immortalised mouse myeloid cells, and an amino-terminal fragment of mouse RECQL4, which contains the regions with homology to yeast Sld2 and worm SLD-2, is sufficient to preserve viability (Buco *et al*., 2025). These data are consistent with past work showing that viable mice can be obtained with deletions within the helicase domain of RECQL4 (Hoki *et al*, 2003; Mann *et al*, 2005), though these mice show defects in skin and skeleton, and have an increased rate of aneuploidy and cancer prevalence. In humans, mutations that truncate RECQL4 within or after the helicase domain are associated with diseases such as Rothmund Thompson syndrome and Rapadilino syndrome, with a range of developmental defects and an increased rate of cancers such as osteosarcoma (Kitao *et al*, 1999; Siitonen *et al*, 2003; Wang *et al*, 2003).

A very recent study showed that MCM10 and RECQL4 also play partially redundant roles during CMG helicase activation in extracts of *Xenopus laevis* eggs (Terui *et al*, 2025), consistent with our findings in worms and mouse cells. Single depletion of RECQL4 was found to inhibit DNA replication *in vitro* to a greater degree than depletion of MCM10, consistent with our findings in worms and mammalian cells.

Finally, it is interesting that CDK phosphorylation of both *C. elegans* SLD-2 (Gaggioli *et al*., 2014) and human RECQL4 (Shin *et al*, 2019; Thakur *et al*, 2025) has been reported to regulate the initiation of DNA replication. Since SLD-2/RECQL4 is dispensable for CMG helicase assembly but required for helicase activation, these findings challenge our view of how CDK controls chromosome replication during DNA replication initiation, suggesting that CDK not only controls assembly of the CMG helicase complex but also regulates its subsequent activation.

## Methods

Resources and reagents from this study are listed in Appendix Table S1 and are available from MRC PPU Reagents and Services (https://mrcppureagents.dundee.ac.uk), or upon request.

### Maintenance of *C. elegans* strains

All *C. elegans* strains in this study are derived from the wild type strain known as ‘Bristol N2’ and are described in Table S1. Alleles generated by genome editing via CRISPR-Cas9 (Suny Biotech) were out-crossed eight times with the N2 wild type before use.

Worms were maintained according to standard procedures (Brenner, 1974) and grown on ‘Nematode Growth Medium’ (NGM: 3 g/l NaCl; 2.5 g/l peptone; 20 g/l agar; 5 mg/l cholesterol; 1 mM CaCl_2_; 1 mM MgSO_4_; 2.7 g/l KH_2_PO_4_; 0.89 g/l K_2_HPO_4_).

### Expression of recombinant C. elegans proteins

To express recombinant worm proteins in budding yeast (see Table S1 for details), the *Saccharomyces cerevisiae* strain yJF1 (*MAT***a** *ade2-1 ura3-1 his3-11,15 trp1-1 leu2-3,112 can1-100 bar1Δ::hphNT pep4Δ::kanMX*) was transformed with the indicated linearised plasmids, containing fragments generated by DNA synthesis (Genscript). Codon usage of the coding sequences for worm proteins was optimised for high-level expression in *Saccharomyces cerevisiae,* as described previously (Yeeles *et al*., 2015).

To expression worm proteins in bacteria, plasmids listed in Table S1 were transformed into the *E. coli* strain Rosetta™ (DE3) pLysS (70956, Novagen).

### Depletion of *C. elegans* proteins by RNA interference

Worms were fed with the RNAse III-deficient bacterial strain HT115 that had been transformed with plasmids expressing double-stranded RNA (plasmids based on the ‘L4440’ parental plasmid - see Table S1). Feeding was performed on 6cm plates that contained the following medium: 3 g/l NaCl, 20 g/l agarose, 5 mg/l cholesterol, 1 mM CaCl_2_, 1 mM MgSO_4_, 2.7 g/l KH_2_PO_4_, 0.89 g/l K_2_HPO_4_, 1 mM IPTG and 100 mg/l Ampicillin.

Plasmids expressing dsRNA were made by cloning PCR products into the vector L4440 (Table S1), after amplification of 1kb cDNA products (*rnr-1*) or full-length cDNA (*cdc-45*, *sld-2*), using a cDNA library that was kindly provided by Sarah-Lena Offenburger and Giulia Saredi. An equivalent plasmid for depletion of SLD-2-GFP was made by amplification of the *GFP-*tag sequence from KAL213. To deplete all four subunits of GINS simultaneously, we cloned contiguous fragments representing the four GINS subunits into a single L4440 plasmid. Details of sequences used in the RNAi vectors are provided in the Table S1. Bacteria expressing *perm-1* dsRNA were obtained from a genome-wide RNAi library (Source Bioscience). Empty L4440 vector was used as the control for the RNAi experiments.

Bacterial cultures were grown to an optical density (OD600) of 1, supplemented with 3 mM IPTG to induce dsRNA expression, and then 400 µl of culture was added to each 6 ml plate, before incubation overnight at room temperature. The next day, 10-20 worms were added to each plate and allowed to feed for 48 hours at 20°C.

### Analysis of *C. elegans* embryos by spinning disk confocal microscopy

Worms at the larval L4 stage were incubated for 48 hours at 20°C on 6 cm plates that had been prepared as above for RNAi feeding. Subsequently, adult worms were dissected in M9 medium (6 g/L Na_2_HPO_4_, 3 g/L KH_2_PO_4_, 5 g/L NaCl, 0.25 g/L MgSO_4_), before five embryos were transferred to a 2% agarose pad and recorded simultaneously from the one-cell stage to the four-cell stage. To monitor events during early S phase of the first embryonic cell cycle, we selected embryos that exhibited mild cortical contractions, which are characteristic of anaphase of the second meiotic division, and then transferred single embryos to a 1 % agarose pad prior to imaging. This procedure was repeated to obtain a total of five embryos for each experimental condition.

Time-lapse video microscopy images were recorded at 24°C with images taken every 10 seconds as described previously (Sonneville *et al*., 2017). We used a Zeiss Cell Observer SD microscope with a Yokogawa CSU-X1 spinning disk and HAMAMATSU C13440 camera, fitted with a PECON incubator and a 60X/1.40 Plan Apochromat oil immersion lens (Olympus). For each time point, a single optical section (z-layer) was captured using ZEN blue software (Zeiss). Subsequently, images were processed and analyzed with ImageJ software (National Institute of Health). The raw images were cropped and the intensity scale adjusted identically for all conditions. The ‘bit depth’ was then changed from 16-bits to 8-bits and videos were assembled. Timepoints were selected and images were further cropped to focus on chromatin. Each image series was then combined into a contiguous sequence, and a Gaussian Blur with a radius of 1 pixel was then applied to all images. Finally, the pixel density was adjusted to 300 dots per inch.

### Monitoring the viability of *C. elegans* embryos

For the experiment in Figure EV1C, experiments were performed in triplicate experiments, using RNAi feeding as above. Subsequently, five adult worms were allowed to produce embryos on a plate during a period of 150 minutes, before the adults were removed and the embryos counted (typically between 60 and 100 embryos). Three days afterwards, the percentage of embryos that had developed into viable adults was determined, together with the mean and standard deviation for each triplicate set.

### EdU incorporation to monitor DNA synthesis in C. elegans embryos

To permeabilize the eggshell to EdU, a bacterial culture expressing *perm-1* dsRNA was diluted 1 in 5 with control bacteria (transformed with the empty L4440 vector), or bacteria expressing GFP dsRNA (for co-depletion of SLD-2-GFP). Otherwise, feeding plates were prepared as described above.

Isotonic growth medium for blastomere culture (Shelton & Bowerman, 1996) was prepared in two steps as follows. Firstly, 1 ml of 5 mg / ml inulin (Sigma-Aldrich, I2255), 50 mg tissue culture–grade polyvinylpyrrolidone (Sigma-Aldrich, P2307), 100 µl Basal Medium Eagle vitamins (Thermo Fisher Scientific, 21010046), 100 µl chemically defined lipid concentrate (Thermo Fisher Scientific, 11905031), 100 µl of 100X concentrated penicillin-streptomycin (Thermo Fisher Scientific, 11568876) and 9 ml Schneider’s Drosophila Medium (Thermo Fisher Scientific, 21720024) were mixed and stored at 4 °C. Secondly, 650µl of this mixture was added to 350 µl of bovine FCS (Thermo Fisher Scientific, A5256701) that had been heat-treated for 30 minutes at 56 °C, giving a final concentration of 35 % FCS.

Permeabilized embryos were then dissected from 10 worms in 150 µl of isotonic growth medium for blastomere culture. After dissection, the mixture of embryos was supplemented with an additional 150 µl of isotonic growth medium containing 40 mM EdU (Thermo Fisher Scientific, E10187), making a final concentration of 20 mM EdU. The embryos were then incubated for 10 minutes, during which time we identified embryos entering the first S phase, on the basis that embryos in early-mid S-phase have strong cortical contractions but have not yet undergone pseudo cleavage. Subsequently, around eight embryos in early-mid S-phase were transferred in 2-3 µl to 1ml fixation solution comprising 4% formaldehyde (Thermo Scientific, 11586711) in Phosphate-buffered saline (PBS) and left at 4 °C for three days.

Finally, embryos were washed twice with PBS, and EdU was detected using the Alexa-647 click-iT EdU Imaging Kit (ThermoFisher Scientific, C10640). After detection, embryos were washed for five minutes in PBS containing 0.05 % Tween-20 (Sigma-aldrich, P1379) and 5 µg/ml Hoechst 33342 (supplied with EdU Imaging Kit), and then washed twice for five minutes in PBS. Embryos were then mounted on 2 % agarose pads and imaged immediately using the Zeiss Cell Observer SD microscope described above. A z-stack of images was acquired from the top to the bottom of each embryo, spanning a depth of ∼ 20 µm in total, with a step size of 0.5 µm. The images shown in Figure 1 represent a maximum-intensity projection of each z-stack.

An ImageJ macro was developed to automate the estimation of EdU intensity. For each z-stack of images corresponding to the female pronucleus (about 15 images per embryo), the EdU signal intensity was calculated as the sum of the Raw Integrated Density within the area encompassing the female pronucleus, after subtraction of background measured from an adjacent cytoplasmic region of identical size.

The experiment was repeated three times to determine mean values and standard deviation. Due to occasional embryo loss during processing, a total of 19–24 embryos were quantified for each condition.

### Purification of *C. elegans* proteins

Proteins purified in this study are listed in the Appendix Table S1. TEV protease (DU6811, MRC PPU Reagents and Services) and PreScission (DU34905, MRC PPU Reagents and Services) protease were kindly provided by Dr. Axel Knebel. Other proteins were produced as described in the following sections, using the following buffer: Buffer A: 25 mM Hepes KOH pH 7.6, 10% glycerol, 0.02% IGEPAL CA-630, 1 mM TCEP.

The following cocktails of protease inhibitors were used as indicated in the sections below: (1) 1X cocktail corresponding to one ‘Roche EDTA free protease inhibitor tablet’ (11873580001, Roche) per 25 ml of buffer (one tablet dissolved in 1 ml water makes a 25X stock solution), together with 0.5 mM PMSF, 5 mM benzamidine HCl, 1 mM AEBSF (A8456, Sigma-Aldrich) and 1 mg/ml Pepstatin A (P5318, Sigma-Aldrich). (2) 1X cocktail corresponding to one Roche EDTA free protease inhibitor tablet (11873580001, Roche) per 25 ml of buffer (one tablet dissolved in 1 ml water makes a 25X stock solution), plus 0.5 mM PMSF.

### Expression of *C. elegans* proteins in budding yeast

The *Saccharomyces cerevisiae* strains used in this study are shown in Appendix Table S1. Yeast cells were grown at 25°C in YP medium (1 % Yeast Extract, 21275, Becton Dickinson; 2 % bacteriological peptone, LP0037B, Oxoid) supplemented with 2 % Raffinose. To express SLD-2 and MUS-101, a 12-litre exponential culture was grown to 1-2 × 10^7^ cells/ml and then synchronized in G1 phase by adding α factor to 0.6 µg/ml for 3.5 hours at 25°C. The G1 culture was induced for 6 hours at 20°C by addition of Galactose to a final concentration of 2 %. To express MCM-10 and RPA, a 12-litre exponential culture was grown to 2-3 × 10^7^ cells/ml and then directly induced for 6 hours at 20°C by addition of Galactose to a final concentration of 2 %. Cells were collected by centrifugation and washed once with lysis buffer (indicated below for each purification) lacking protease inhibitors. Cell pellets (∼ 30 g) were then resuspended in 0.3 volumes of the indicated lysis buffer containing protease inhibitors. The resulting suspensions were then frozen dropwise in liquid nitrogen and stored at −80°C. Subsequently, the entire sample of frozen yeast cells were ground in the presence of liquid nitrogen, using a SPEX CertiPrep 6850 Freezer/Mill with 3 cycles of 2 minutes at a rate of 15. The resulting powders were then stored at −80 °C.

The CMG helicase was purified as previously described (Xia *et al*., 2023). TIM-1_TIPN-1, CTF-4, CLSP-1,and CTF-18_RFC were purified as previously described (Xia *et al*., 2021).

#### C.e. MUS-101

Yeast cell powder was thawed in buffer A / 0.5 M NaCl / 1X protease inhibitor cocktail 1. The mixture was centrifuged at 100,000 x g at 4°C for 0.5 hour, followed by another step of centrifugation at 235,000 x g at 4°C for 1 hour. After spinning, the soluble extract was recovered and mixed with 3 ml IgG resin (17096901, GE). The mixture was incubated at 4 °C for 2 hours with rotation.

Resin was collected and washed extensively with buffer A / 0.5 M NaCl / 1X protease inhibitor cocktail 1. The resin was then incubated with 40ml buffer A / 0.5 M NaCl / 10 mM Mg(OAc)_2_ / 2 mM ATP at 4 °C for 10 minutes to remove chaperones, then washed extensively with buffer A / 0.5 M NaCl. The purified proteins were then eluted by overnight incubation with rotation in 2 ml buffer A / 0.5 M NaCl containing 100 µg TEV protease.

The supernatant was collected, and the resin was further eluted twice with 4ml buffer A / 0.5 M NaCl. The elute was concentrated and loaded onto a 24 ml Superdex 200 column in buffer A / 0.5 M NaCl. Peak fractions containing MUS-101 were pooled, aliquoted, and snap frozen in liquid nitrogen and stored at – 80 °C.

#### C.e. SLD-2

Yeast cell powder was thawed in buffer A / 0.5 M NaCl / 1X protease inhibitor cocktail 1. The mixture was centrifuged at 100,000 x g at 4°C for 0.5 hours, followed by another step of centrifugation at 235,000 x g at 4°C for 1 hours. After spinning, the soluble extract was recovered and mixed with 3 ml IgG resin (17096901, GE). The mixture was incubated at 4 °C for 2 hours with rotation.

Resin was collected and washed extensively with buffer A / 0.5 M NaCl / 1X protease inhibitor cocktail 1. The resin was then incubated with 40ml buffer A / 0.5 M NaCl / 10 mM Mg(OAc)_2_ / 2 mM ATP at 4 °C for 10 minutes to remove chaperones, then washed extensively with buffer A / 0.5 M NaCl. The purified proteins were then eluted by 20 hours incubation with rotation in 2 ml buffer A / 0.5 M NaCl containing 100 µg TEV protease.

The supernatant was collected, and the resin was further eluted twice with 2ml buffer A / 0.5 M NaCl. The elute was concentrated and loaded onto a 24 ml Superdex 200 column in buffer A / 0.5 M NaCl. Peak fractions containing SLD-2 were pooled, concentrated and re-loaded onto a 24 ml Superdex 200 column in buffer A / 0.5 M NaCl. Peak fractions were pooled, aliquoted, and snap frozen in liquid nitrogen and stored at −80 °C.

#### MCM-10

Yeast cell powder was thawed in buffer A / 0.2 M NaCl / 1X protease inhibitor cocktail 1. The mixture was centrifuged at 100,000 x g at 4°C for 0.5 hours, followed by another step of centrifugation at 235,000 x g at 4°C for 1 hour. After spinning, the soluble extract was recovered and mixed with 3 ml IgG resin (17096901, GE). The mixture was incubated at 4 °C for 2 hours with rotation.

Resin was collected and washed extensively with buffer A / 0.2 M NaCl / 1X protease inhibitor cocktail 1. The resin was then incubated with 40ml buffer A / 0.2 M NaCl / 10 mM Mg(OAc)_2_ / 2 mM ATP at 4 °C for 10 minutes to remove chaperones, then washed extensively with buffer A / 0.2 M NaCl. The purified proteins were then eluted by overnight incubation with rotation in 2 ml buffer A / 0.2 M NaCl containing 100 µg TEV protease.

The supernatant was collected, and the resin was further eluted up to 10ml with buffer A / 0.2 M NaCl. The eluate was loaded onto a 1 ml HiTrap SP HP column in buffer A / 0.2 M NaCl. MCM-10 was eluted with a 20 ml gradient from 0.2 – 0.6 M NaCl. The peak fractions were pooled, concentrated, aliquoted, snap frozen in liquid nitrogen and stored at −80 °C.

#### C.e. RPA

Yeast cell powder was thawed in buffer A / 0.5 M NaCl / 1X protease inhibitor cocktail 1. The mixture was centrifuged at 100,000 x g at 4°C for 0.5 hours, followed by another step of centrifugation at 235,000 x g at 4°C for 1 hour. After spinning, the soluble extract was recovered and mixed with 3 ml IgG resin (17096901, GE). The mixture was incubated at 4 °C for 2 hours with rotation.

Resin was collected and washed extensively with buffer A / 0.5 M NaCl / 1X protease inhibitor cocktail 1. The resin was then incubated with 40ml buffer A / 0.5 M NaCl / 10 mM Mg(OAc)_2_ / 2 mM ATP at 4 °C for 10 minutes to remove chaperones, then washed extensively with buffer A / 0.5 M NaCl. The purified proteins were then eluted by overnight incubation with rotation in 2 ml buffer A / 0.5 M NaCl containing 100 µg TEV protease.

The supernatant was collected, and the resin was further eluted twice with 4ml buffer A / 0.5 M NaCl. The elute was concentrated and loaded onto a 24 ml Superdex 200 column in buffer A / 0.5 M NaCl. Peak fractions containing RPA were pooled, aliquoted, and snap frozen in liquid nitrogen and stored at −80 °C.

### Protein expression in *E. coli*

*C. elegans* DNSN-1 and the truncated or mutated variants DNSN-1-ΔN and DNSN-1-3A were purified as previously described (Xia *et al*., 2023).

### Preparation of DNA substrates for helicase and DNA binding assays

For the DNA helicase assays in Figure 4 and Figure EV3, model DNA replication forks were assembled *in vitro* from annealed oligonucleotides. Firstly, 400 nM of PAGE-purified 5’ oligonucleotides (see Appendix Table S1 for oligonucleotide sequences) were labeled with γ-[32P]-ATP using 10 Units of T4 Polynucleotide Kinase (T4 PNK, New England Biolabs M0201S), in a 10 µl volume for 30 minutes at 37°C. The reactions were then heated at 95 °C for 10 minutes to inactive T4 PNK. The 10 µl labeling reaction was then mixed with 200 nM 3’ oligonucleotides (see Appendix Table S1 for oligonucleotide sequences), 100 mM NaCl, and 10 mM Mg(OAc)_2_ in a total volume of 20 µl, before heating to 95 °C for 10 minutes in a metal heating block. Finally, the metal block was placed at room temperature and the sample left to cool for 3 h, allowing the oligonucleotides to anneal with each other. Unincorporated γ-[32P]-ATP was then removed using Cytiva MicroSpin G-50 columns (ThermoFisher Scientific, 27533001).

To prepare a double-stranded DNA substrate for the DNA binding assay in Figure EV4, 500 nM of Cy5-labeled 80 nt 5’ oligonucleotides was incubated with 500nM of 80 nt 3’ oligonucleotides (see Appendix Table S1 for oligonucleotide sequences), in a total volume of 20 µl containing 100 mM NaCl and 10 mM Mg(OAc)_2_. The reactions were then heated to 95 °C for 10 minutes before cooling to room temperature, as described above.

### CMG helicase assays

CMG helicase assays were conducted in a ‘helicase buffer’ consisting of 25 mM HEPES-KOH (pH 7.6), 60 mM potassium glutamate, 0.02% NP-40-S, 1 mM DTT, 10 mM Mg(OAc)2, 0.1 mM EDTA and 0.2 mg/ml Bovine Serum Albumin. Unless indicated otherwise in the figure legends, the DNA, protein and nucleotide concentrations in the final reaction were: 1 nM fork DNA, 30 nM CMG, 120 nM MCM-10, DNSN-1, or SLD-2, 5 mM ATP, plus 40 nM unlabelled 5’ oligonucleotides as ‘trap DNA’ to prevent re-annealing of the product. Reactions were set up as follows: recombinant *C. elegans* CMG (30 nM) was pre-incubated with the DNA substrate (1 nM) at 25 °C for 15 minutes in helicase buffer supplemented with 0.1 mM AMP-PNP. Reactions were then initiated by addition of a 10X solution containing ATP, trap DNA and MCM10, DNSN-1 or SLD-2, before incubation at 25 °C for 30 minutes. The reactions were then stopped by addition of 3 µl of buffer containing 250 mM EDTA, 5% SDS and 0.6 Units of proteinase K, and the incubation continued at 37 °C for 1 hour. The samples were supplemented with Novex Hi-Density TBE Sample Buffer (ThermoFisher Scientific, LC6678) and analyzed in 4%–20% Novex TBE Gels (ThermoFisher Scientific, EC62252BOX) at 200 V for 30 minutes in 0.5X TBE with 0.1% SDS. Gels were mounted onto chromatography paper (GE Healthcare, 3030-861) and then exposed to BAS-MS Imaging Plates (Fujifilm), which were developed on a Typhoon phosphorimager (GE Healthcare).

### DNA binding assay

DNA binding reactions were set up as follows: 15 nM, 45 nM, or 135 nM of indicated protein was pre-incubated with 5 nM Cy5-labelled DNA at 25 °C for 15 minutes, in ‘DNA binding buffer’ consisting of 25 mM HEPES-KOH (pH 7.6), 50 mM potassium glutamate, 0.02% NP-40-S, 1 mM DTT, 10 mM Mg(OAc)2 and 10% glycerol. The samples were analyzed in 4%–20% Novex TBE Gels (ThermoFisher Scientific, EC62252BOX) at 200 V for 40 minutes in 0.5X TBE. The gels were then analyzed with a Bio-Rad ChemiDoc.

### Immunoprecipitation of reconstituted complexes of *C. elegans* proteins

Reactions with a final volume of 10 µl were set up on ice as follows: 25 mM Hepes-KOH (pH 7.6), 0.02% IGEPAL CA-630, 0.1 mg / ml BSA, 1 mM DTT, 100 mM KOAc, 10 mM Mg(OAc)_2_, 0.5 mM AMP-PNP, 50nM DNA substrate comprising 46bp double-stranded DNA and a 39 nt ‘3’-flap’ of single-stranded DNA (sequences are shown in Appendix Table S1) and 3.3 µl ‘protein mix’ containing 15 nM CMG and either MCM-10 (30 nM), SLD-2 (30 nM), or DNSN-1 (30 nM dimer), in 300 mM KOAc (giving a final concentration of 200 mM KOAc). Reactions were incubated on ice for 30 minutes, before addition of 5 µl magnetic beads (Dynabeads M-270 Epoxy; 14302D, ThermoFisher Scientific) that had been coupled to anti-MCM-6 antibodies as described below, in order to bind the CMG helicase complex. After 1 hour, protein complexes bound to the magnetic beads were washed twice with 1 ml of buffer containing 25 mM Hepes-KOH (pH 7.6), 0.02% IGEPAL CA-630, 0.1 mg / ml BSA, 1 mM DTT, 10 mM Mg(OAc)_2_ and 200 mM KOAc. The bound proteins were eluted at 95°C for 5 minutes in 30 µl of 1X Laemmli buffer.

### Culture of mouse ES cells

Mouse ES cells of the E14tg2A background were maintained at 37°C in a humidified atmosphere of 5% CO_2_, 95% air in Dulbecco’s Modified Eagle Medium (DMEM, ThermoFisher Scientific, 11960044) supplemented with 10% Foetal Bovine Serum (FBS, FCS-SA/500, LabTech), 5% KnockOut Serum Replacement (ThermoFisher Scientific, 10828028), 2 mM L-Glutamine (ThermoFisher Scientific, 25030081), 100 U/ml Penicillin-Streptomycin (ThermoFisher Scientific, 15140122), 1 mM Sodium Pyruvate (ThermoFisher Scientific, 11360070), 1X non-essential amino acids (ThermoFisher Scientific, 11140050), 0.05 mM β-mercaptoethanol (Sigma-Aldrich, M6250) and 0.1 μg/ml LIF (MRC PPU Reagents and Services, DU1715). Around 2 × 10^6^ cells were plated on 10 cm plates precoated with 0.1% gelatin (Sigma, G1890), and then on alternate days the cells were released using 0.05 % Trypsin-EDTA (ThermoFisher Scientific, 25300054), before 1 in 10 dilution in fresh medium and re-plating.

### CRISPR-Cas9 genome editing, transfection of mouse ES cells and selection of clones

We used the ‘CRISPR Finder’ provided by the Welcome Sanger Institute (https://wge.stemcell.sanger.ac.uk//find_crisprs) to design pairs of guide RNAs (gRNAs) to target each specific site of interest in the mouse genome. For each gRNA, two oligos containing the homology region to the targeted gene were annealed and cloned into the vectors pX335 or pKN7 after digestion by BbsI, as previously described previously (Pyzocha *et al*, 2014).

To introduce tags at the amino termini of RECQL4 or MCM10, cells were transfected with 2 µg each of three plasmids. The first plasmid expressed the Cas9-D10A nickase and a gRNA, the second expressed the Puromycin resistance gene and a second gRNA. The third was a ‘donor plasmid’ in which the chosen tag was flanked by around 500-1000 bp homology to the targeted gene, together with the Hygromycin or Blasticidin resistance gene. The plasmids were added to 200 µl of DMEM plus 15 µl of 1 mg/ml linear Polyethylenimine (PEI; Polysciences, Inc, 24765-2). Subsequently, 1.0 × 10^6^ cells were gently mixed with the DNA-PEI-DMEM mix and incubated at room temperature for 30 minutes. Cells were then transferred to a single well of a 6-well plate containing 2 ml of complete DMEM. Cells were subjected to two 24-hour rounds of selection with fresh medium containing 2 μg/ml Puromycin (ThermoFisher Scientific, A1113802) followed by one to two weeks of selection with fresh medium containing 200-500 µg/ml Hygromycin (Invitrogen, ant-hg-5) or 7.5 μg/ml Blasticidin (Invitrogen, ant-bl-10p) as appropriate, to select for the marker in the donor vector (Hygromycin-resistance for *BromoTag-Mcm10* and Blasticidin-resistance for *dTAG-Recql4*). A series of dilutions from 10 to 10,000 times was made of the surviving cells, in complete DMEM containing antibiotic corresponding to the second selection marker, before replating to allow the picking of single colonies after five to seven days additional incubation. Cells were re-plated once again in selective medium, and then after six to ten days the clones were expanded and subsequently analysed by immunoblotting, PCR and DNA sequencing of the target locus.

To insert the GFP tag into the *Psf1* locus in *BromoTag-Mcm10 dTAG-Recql4* cells, the procedure was similar except that the donor plasmid lacked a selective marker, and cells were initially subjected to two 24-hour rounds of Puromycin selection. Subsequently, viable GFP-positive cells were sorted by flow cytometry into individual wells of a 96-well plate containing complete DMEM lacking antibiotics.

To create small deletions in exon 1 of *Recql4* or exon 3 of *Mcm10*, cells were transfected with plasmids expressing the Cas9-D10A nickase, two gRNAs and Puromycin resistance. After two 24-hour rounds of Puromycin selection, surviving cells were sorted by flow cytometry into individual wells of a 96-well plate containing complete DMEM.

To delete the entire *Mcm10* or *Recql4* gene, as summarised in Figures EV7-8, we adapted a previously described protocol (Saito *et al*, 2024). Cells were transfected with plasmids expressing wild-type Cas9, two gRNAs targeting the 5’ and 3’ end of *Mcm10* or *Recql4*, and the Puromycin-resistance gene. After two 24-hour rounds of Puromycin selection, surviving cells were diluted and then 50-100 cells were plated onto new 10 cm plates and incubated until colonies formed. Single colonies were then picked and grown in a 24-well plates containing complete DMEM lacking antibiotics.

To express *Recql4* from the *Rosa26* ‘safe haven’ locus, we transfected cells with two plasmids. The first expressed the Cas9-D10A nickase, the Puromycin-resistance gene, and a single gRNA for *Rosa26* as described previously (Villa *et al*., 2021). The second was a donor plasmid expressing wild type or mutant RECQL4 from the constitutive EF1a promotor, together with a Neomycin resistance gene.

These sequences in the donor vector were flanked by around 800 bp 5’ and 3’ homology to the *Rosa26* locus. After two rounds of 24-hour selection with puromycin, integration of the donor DNA was selected by exposing cells for one to two weeks to fresh medium containing 700 μg/ml G418 (Formedium, G418S). Single clones were then selected and analysed as described above for *BromoTag-Mcm10* and *dTAG-Recql4*.

### PCR and sequence analysis for genotyping of mouse ES cells

Cells from a 10cm plate were released as described above. One third of the cells were pelleted and resuspended in 100 μl of 50 mM NaOH and the sample was heated at 95°C for 15 minutes. The pH was then neutralised by addition of 11 µl of 1 M Tris-HCl (pH 6.9). Subsequently, a PCR reaction was performed with PrimeStar Hot Start DNA polymerase (Takara Bio, R010A) and the product was subcloned with StrataClone Blunt PCR cloning kit (Agilent Technologies, 240207), before DNA sequenced with M13 Forward, M13 Reverse primers and primers specific to the target locus (see Appendix Table S1).

### Cell proliferation assay for mouse ES cells

In Figure 5A-B, cells were transfected as indicated with pairs of gRNAs to create small deletions at the beginning of exon 1 of *Recql4* or exon 3 of Mcm10, as described above (plasmids expressing gRNAs and the Cas9-D10A nickase were omitted in the control). After two rounds of puromycin selection, 500 cells (*Control* and *Mcm10Δ*) or 1000 cells (*Recql4Δ*) were seeded on a 10 cm dish and incubated at 37 °C for seven days (*Control* and *Mcm10Δ*) or 10 days (Recql4Δ). The cells were then washed with Phosphate-buffered saline (PBS) and fixed with 20 % methanol. Subsequently, cells were stained with 0.5 % crystal violet solution (Sigma-Aldrich, HT90132) and the plates were scanned.

In Figure 5C, 500 cells were seeded on a 10 cm dish. After 24 hours, 250 nM AGB1 and 250 nM dTAG13 were added to the medium, before incubation at 37 °C for seven days. The cells were then washed with PBS and fixed with 20 % methanol, before staining and scanning as above.

Generation times were measured in at least three independent experiments, before calculation of mean and standard deviation.

### Denatured protein extracts of mouse ES cells

Firstly, 1.0 × 10^6^ cells were seeded in a 6-well plate. The next day, the cells were treated with 250 nM AGB1(Tocris, 7686) and / or 250 nM dTAG13 for 3 hours, as indicated in Figure 5D and Figure 7A+F. Cells were then released from the dish with 0.05% Trypsin-EDTA and washed with PBS. The pellet was resuspended in 150 µl of 20% TCA and then 150 µl of acid-washed glass beads (Stratech, 11079105) was added, before vortexing for one minute. The supernatant was then transferred into a fresh 1.5 ml Eppendorf tube. The glass beads were then washed with 150 µl of 5% TCA, and this second supernatant was added to the first one, before centrifugation for 10 minutes at 845 x g at room temperature. The pellet was then resuspended in 37.5 µl ‘High pH Laemmli buffer’ (150 µl of 1M Tris base per 1ml of 1X Laemmli buffer) and 12.5 µl 4xNuPAGE LDS sample buffer (ThermoFisher Scientific, NP0007) before incubation at 95 °C for five minutes. For the experiment in Figure 8E, 250 nM PROTACs or 2 mM Hydroxyurea (HU, Molekula, 10872383) was added to the cells for four or eight hours as indicated.

### Synchronisation of mouse ES cells with respect to the cell cycle

For the experiment in Figure 6, 2.5 × 10^6^ cells were seeded on a 10 cm dish for subsequent flow cytometry, or 5 × 10^6^ cells on a 20 cm dish for the generation of native cell extracts for immunoprecipitation analysis. After 24 hours, 1.25 mM thymidine (Sigma-Aldrich, T9250) was added to cells for 18 hours to synchronise them in early S- phase. The cells were then washed twice with PBS and then incubated for nine hours in fresh medium containing 5 µM S-trityl-L-cysteine or STLC (Sigma-Aldrich, 164739), which inhibits the kinesin Eg5 and arrests cells in mitosis. During the last two hours of the arrest with STLC, 250 nM of PROTACs (both AGB1 and dTAG13) was added to degrade both BromoTag-MCM10 and dTAG-RECQL4. The cells were then washed three times with PBS and released into fresh medium in the presence or absence of 250 nM PROTACs as indicated in Figure 6.

### Measuring DNA content of mouse ES cells by flow cytometry

Cells were released from a 10 cm dish and fixed with 1 ml 70 % ethanol. Subsequently, 300 μl of fixed cells were washed once with PBS and then resuspended in 1 ml PBS containing 0.05 mg/ml propidium iodide (Sigma-Aldrich, P44170) and 0.05 mg/ml RNase A (ThermoFisher Scientific, EN0531). The cells were then incubated at 24°C for one hour to digest RNA. The samples were processed with an LRS Fortessa flow cytometer (Becton Dickinson) and the data analysed with FlowJo software.

### Immunoprecipitation of protein complexes from extracts of mouse ES cells

For the experiment in Figure 6C, four 15 cm dishes were used for each sample. Cells were released from the plates by incubation with PBS containing 1mM EDTA and 1mM EGTA for 10 minutes and then harvested by centrifugation. The cells were then resuspended in one volume of lysis buffer: 100 mM Hepes-KOH pH 7.9, 100 mM potassium acetate, 10 mM magnesium acetate, 2 mM EDTA, 10% glycerol, 0.1% Triton X-100, 2 mM sodium fluoride, 2 mM sodium β-glycerophosphate pentahydrate, 1 mM dithiothreitol, 1% Protease Inhibitor Cocktail (Sigma-Aldrich, P8215), 1x Complete Protease Inhibitor (Roche, 05056489001). Chromosomal DNA was digested for 30 minutes at 4°C by addition of 1600 U/ml of Pierce Universal Nuclease (ThermoFisher Scientific, 88702), before centrifugation at 20,000 x g for 30 minutes at 4°C. The resulting extract was added to 20 µl ChromoTekGFP-Trap magnetic Particles M-270 (ThermoFisher Scientific, 17353353) and incubated for two hours at 4°C. Bound proteins were eluted by heating the beads at 95°C for five minutes in presence of 50 µl Laemmli buffer. Proteins were separated on a 4-12% Bis-Tris NuPAGE gel (Life Technologies, NP0301) in 1X MOPS buffer, and then detected by immunoblotting.

### High-content screening microscopy to monitor CMG assembly in mouse ES cells

For the experiments in Figures 7-8, and Figure EV10, mouse ES cells were seeded at 20,000 cells per well in clear-bottomed black wall 96-well plates (Greiner, 655090). Cells were grown for 24 hours, followed by an additional eight hours in the presence or absence as indicated in the figures of 250 nM AGB1, 250 nM dTAG13, or 2 mM hydroxyurea. Cells were then treated with 20 μM EdU for 20 minutes, except for a small number of wells that were left untreated to serve as controls for the detection of EdU incorporation. Cells were then washed with cold PBS containing 1 mM CaCl_2_, 1 mM MgCl_2_, and 20 µM thymidine. To remove proteins not bound to chromatin, cells were pre-extracted for 5 minutes with cold pre-extraction buffer (20 mM HEPES-KOH pH 7.6, 20 mM NaCl, 5 mM MgCl_2_, 0.5 % NP40, 1 mM DTT) supplemented with protease inhibitors (Roche, Complete EDTA-free, 11873580001), before fixation 15 minutes in 3.5 % formaldehyde in PBS. Cells were then washed with PBS containing 1 mM CaCl_2_, 1 mM MgCl_2_. Subsequently, EdU was covalently linked to the fluorescent dye Alexa-647 using the click-iT EdU Imaging Kit (ThermoFisher Scientific, B10184). Cells were then washed in PBS followed by incubation in methanol for 15 minutes at 4 °C and then blocked for 30 minutes with 5 % BSA in PBS supplemented with 0.05 % Tween 20, before incubation with a 1 in 50 dilution of an anti-CDC45 antibody (Cell Signalling Technology, 11881, rabbit monoclonal) for 66 hours at 4 °C. Cells were then incubated with a 1 in 250 dilution of anti-MCM2 antibody (BD Biosciences, 610701, mouse monoclonal) for three hours at room temperature in the dark. Detection of RPA32 S4 / S8 phosphorylation was performed on cells without anti-CDC45 incubation by incubating cells with a 1 in 200 dilution of Phospho-RPA32 (Ser4, Ser8) antibody (Thermofisher/Bethyl, A300-245A, rabbit polyclonal) overnight at 4 °C. Subsequently, cells were washed in 5 % BSA / PBST followed by incubation with donkey anti-rabbit IgG coupled to Alexa Fluor 488 (Invitrogen, A-21206) or goat anti-mouse IgG coupled to Alexa Fluor 546 (Invitrogen, A-11030) for 1 hour at room temperature, in the presence of 2.5 μg/ml 4′,6-diamidino-2-phenylindole or DAPI (Thermo, 62248). For detection of GFP-PSF1, cells were prepared as above but the methanol fixation and antibody incubation steps were omitted.

Finally, cells were washed and stored in PBS at 4°C until imaging with the scanR High-Content Screening Microscope (Olympus). Images were analysed with the ScanR analysis software and data visualised in Tableau. For CDC45 and MCM2 plots in Figure 7C-I, Figure 8B-C and Figure EV10B-C, the EdU-negative population of cells is coloured black (the threshold was established using the intensity of the EdU non-treated control wells and then applied to the rest of the plate).

### Live-cell imaging of GFP-PSF1 on heterochromatin patches in mouse ES cells

For the experiment in Figure 8F, live mouse ES cells were stained with a 4000 fold dilution of SPY650-DNA (SC501, Spirochrome), before imaging by spinning-disk confocal microscopy as described above for *C. elegans*. Over 600 cells were examined for each sample, and the proportion of cells in which GFP-PSF1 co-localised with heterochromatin patches was determined.

## Acknowledgements

We thank Qasim Ashraf for assisting RS with initial studies of *C. elegans* SLD-2, Lucia Puchades Gimeno for helping CE with deletion of the *Recql4* gene, and MRC PPU Reagents and Services (https://mrcppureagents.dundee.ac.uk) for antibody production including DNA cloning. We gratefully acknowledge the support of Cancer Research UK (Discovery Programme Award DRCRPG-Nov22/100016 to KL) and Wellcome (Discovery Award 302391/Z/23/Z to KL). Materials generated in this study are listed in Appendix Table S1 and are available from MRC PPU Reagents and Services (https://mrcppureagents.dundee.ac.uk) or upon request.

---

**Figure EV1.**
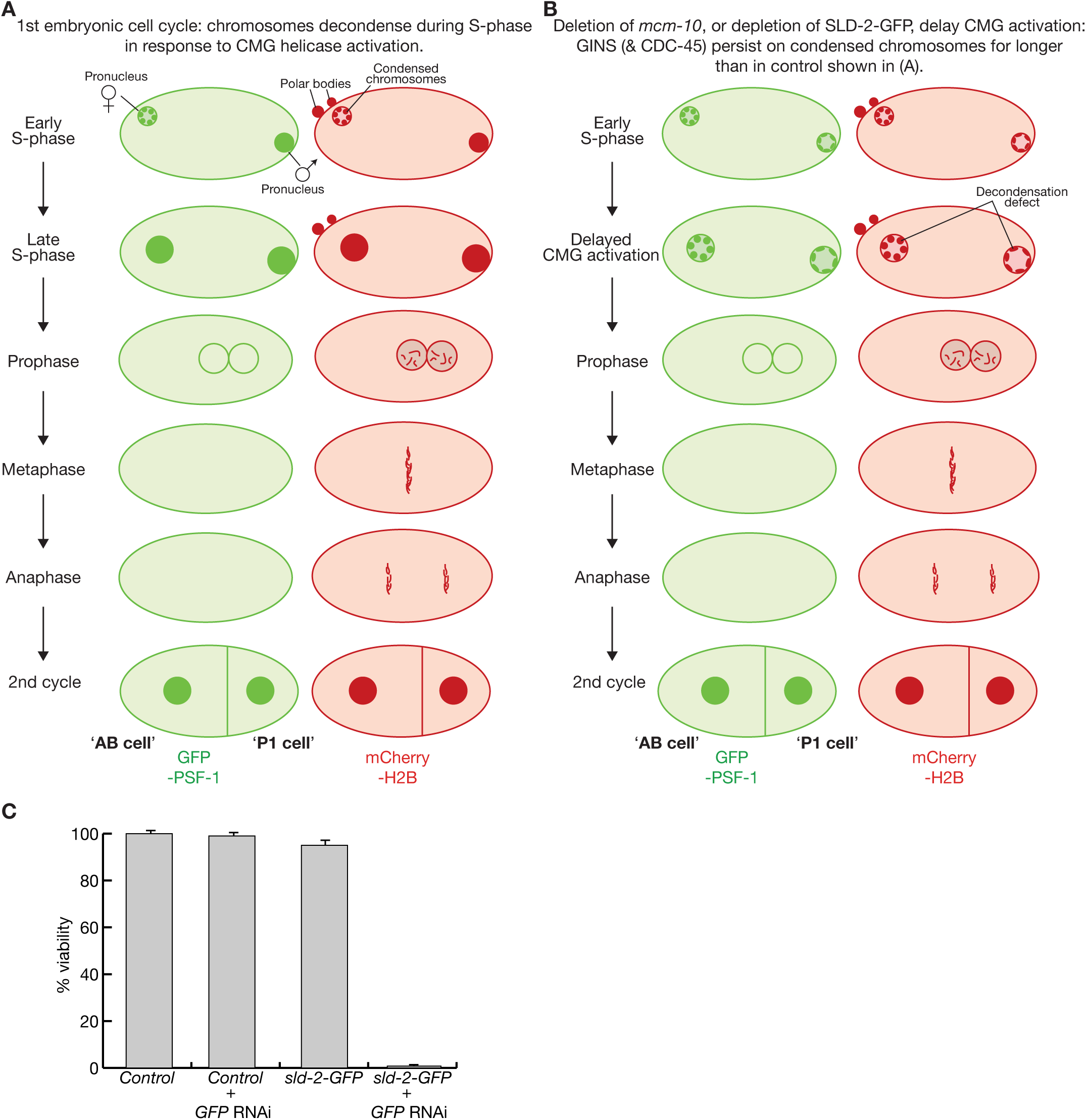
Depletion of SLD-2 is lethal in *C. elegans*. (**A**) Illustration of the first embryonic cell cycle of C. elegans expressing GFP-PSF-1 and mCherry-HistoneH2B. (**B**) Illustration of how chromosome decondensation is delayed in response to defects in CMG helicase activation in the first embryonic cell cycle of *mcm-10Δ* worms, or in response to depletion of SLD-2. (**C**) Embryonic viability was measured as described in Methods, after treating control (wild type ‘N2’ worms) or *sld-2-GFP* worms with anti-GFP RNAi as indicated. Mean values and standard deviations are presented from three independent experiments.

**Figure EV2.**
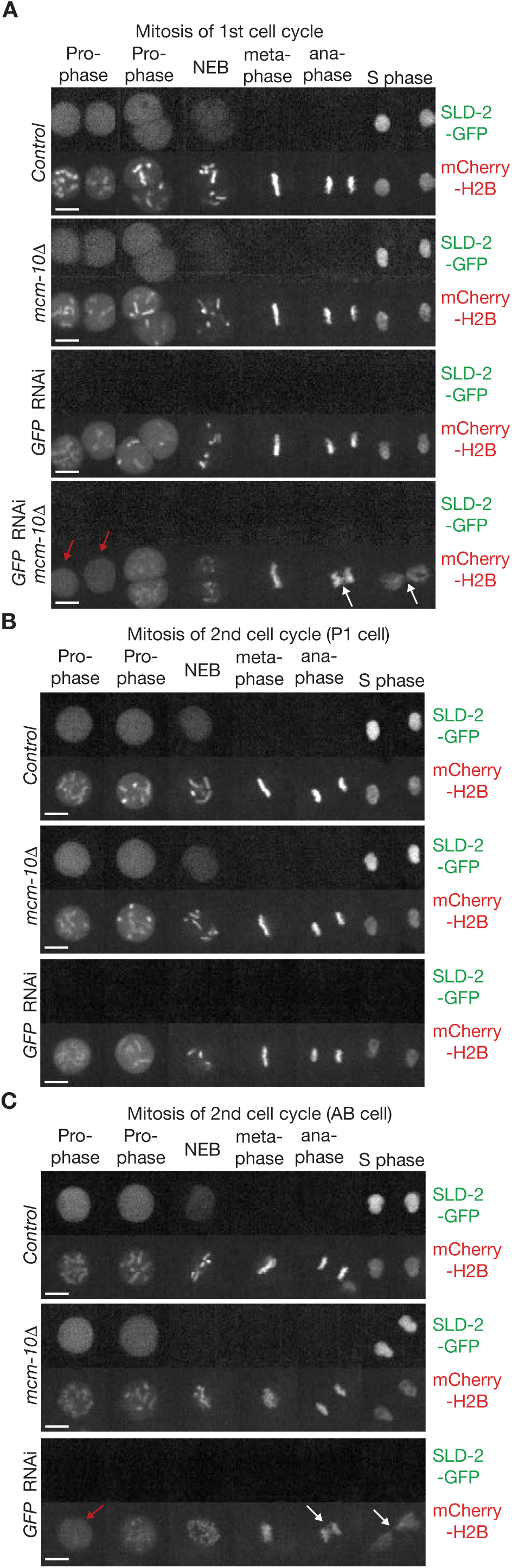
Mitosis proceeds normally in *mcm-10Δ* worms but fails during the second cell cycle in the AB cell of worms lacking SLD-2. (**A**) Progression through mitosis of the first embryonic cell cycle was monitored by time-lapse video microscopy of control and *mcm-10Δ* worms expressing SLD-2-GFP and mCherry-HistoneH2B, after treatment as indicated with anti-GFP RNAi. Red arrows highlight defective chromosome condensation during prophase, whereas white arrows indicate the failure of chromosome segregation. Both defects were observed in all *mcm-10Δ sld-2-GFP* embryos from worms that had been treated with GFP RNAi (n = 5). (**B**) Progression of the P1 cell through mitosis of the second embryonic cell cycle, in the same experiment as in (A). (**C**) Progression of the AB cell through mitosis of the second embryonic cell cycle, in the same experiment as in (A). The red arrow indicates the defect in mitotic chromosome condensation in prophase, whereas white arrows denote the failure of chromosome segregation during anaphase. Both defects were observed in all *sld-2-GFP* embryos from worms treated with GFP RNAi (n = 5), indicating that cells had entered mitosis before completion of chromosome replication.

**Figure EV3.**
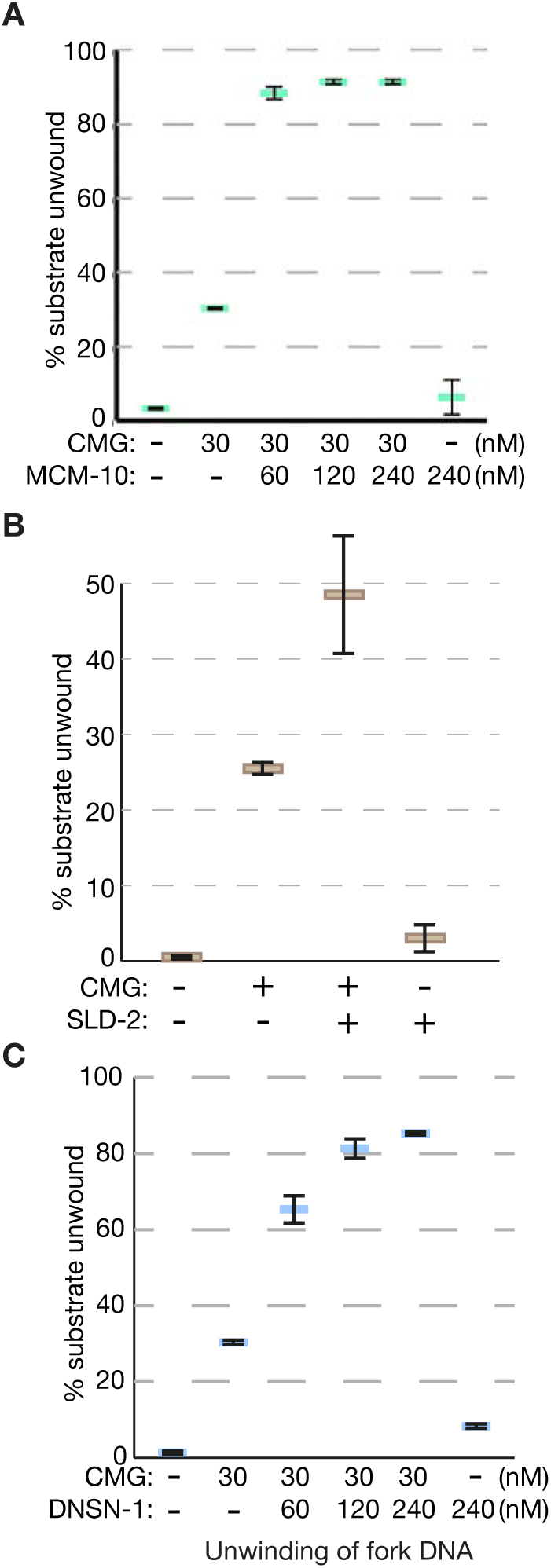
Stimulation of CMG helicase activity by MCM-10, SLD-2 and DNSN-1. (**A**) CMG helicase activity was monitored as for Figure 4 and as described in Methods, in the presence of the indicated concentrations of CMG and MCM-10. The data correspond to the mean values and standard deviation from three independent experiments. (**B**) Similar experiment in the presence of the indicated concentrations of CMG and SLD-2. (**C**) Analogous experiment in the presence of the indicated concentrations of CMG and DNSN-1.

**Figure EV4.**
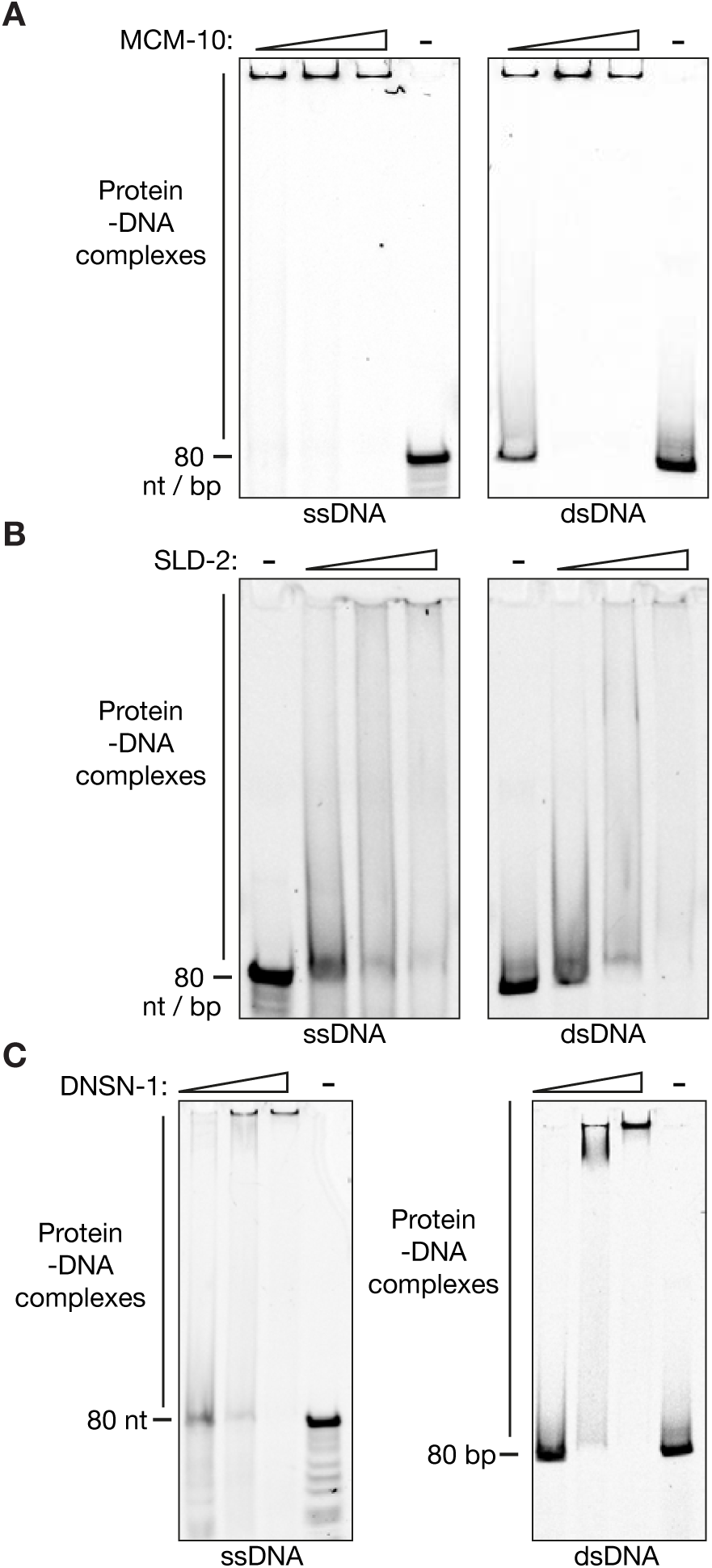
MCM-10, SLD-2 and DNSN-1 all bind to DNA *in vitro*. As described in Methods, the indicated concentrations of MCM-10 (A), SLD-2 (B) and DNSN-1 (C) were incubated with 5 nM Cy5-labelled DNA at 25 °C for 15 minutes. Protein-DNA complexes were then analyzed in native polyacrylamide gradient gels before detection of the fluorescently-labelled DNA.

**Figure EV5.**
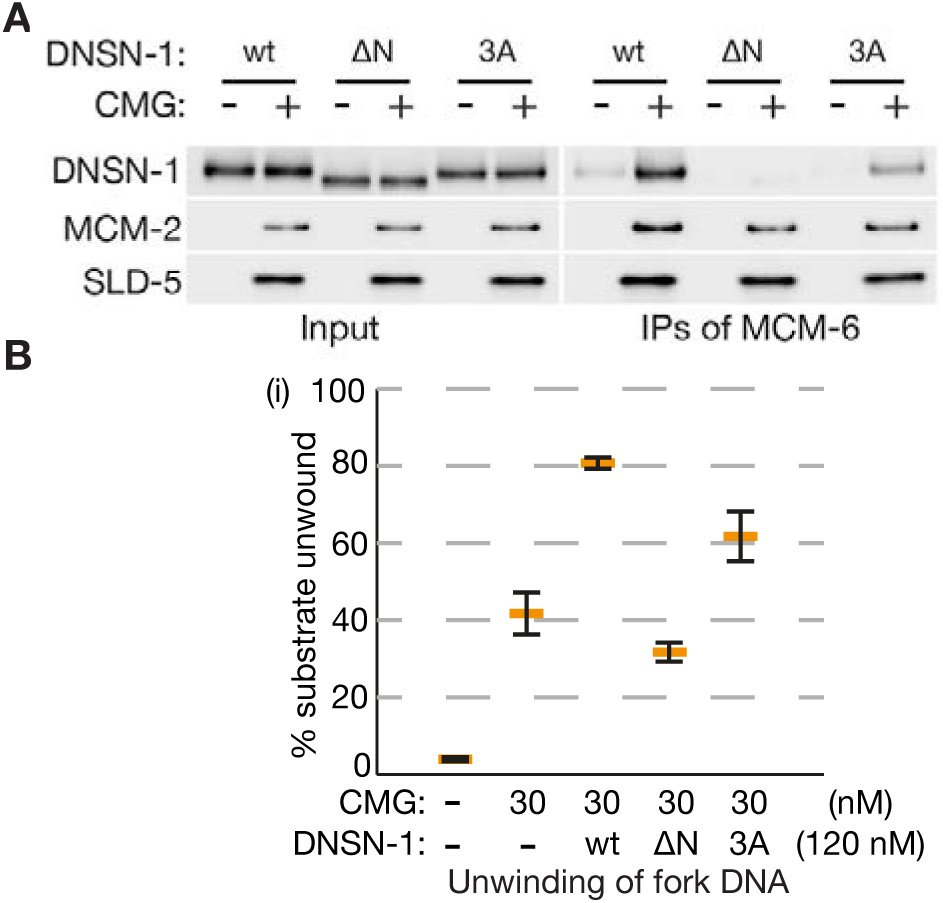
The ability of DNSN-1 to bind CMG is important for the stimulation of helicase activity *in vitro*. (**A**) Recombinant *C. elegans* CMG was mixed as indicated with wild type DNSN-1 (wt), DNSN-1 lacking the first 20 amino acids (DNSN-1-ΔN), or DNSN-1 with mutations in three conserved residues that contact the MCM-3 subunit of the CMG helicase (DNSN-1-3A). As described above for Figure 2C and 2F, the CMG helicase was then isolated by immunoprecipitation of the MCM-6 subunit, and the indicated proteins detected by immunoblotting. (**B**) The ability of DNSN-1, DNSN-1-ΔN and DNSN-1-3A to stimulate CMG helicase activity was monitored in vitro as for Figure 4C and Figure EV3C.

**Figure EV6.**
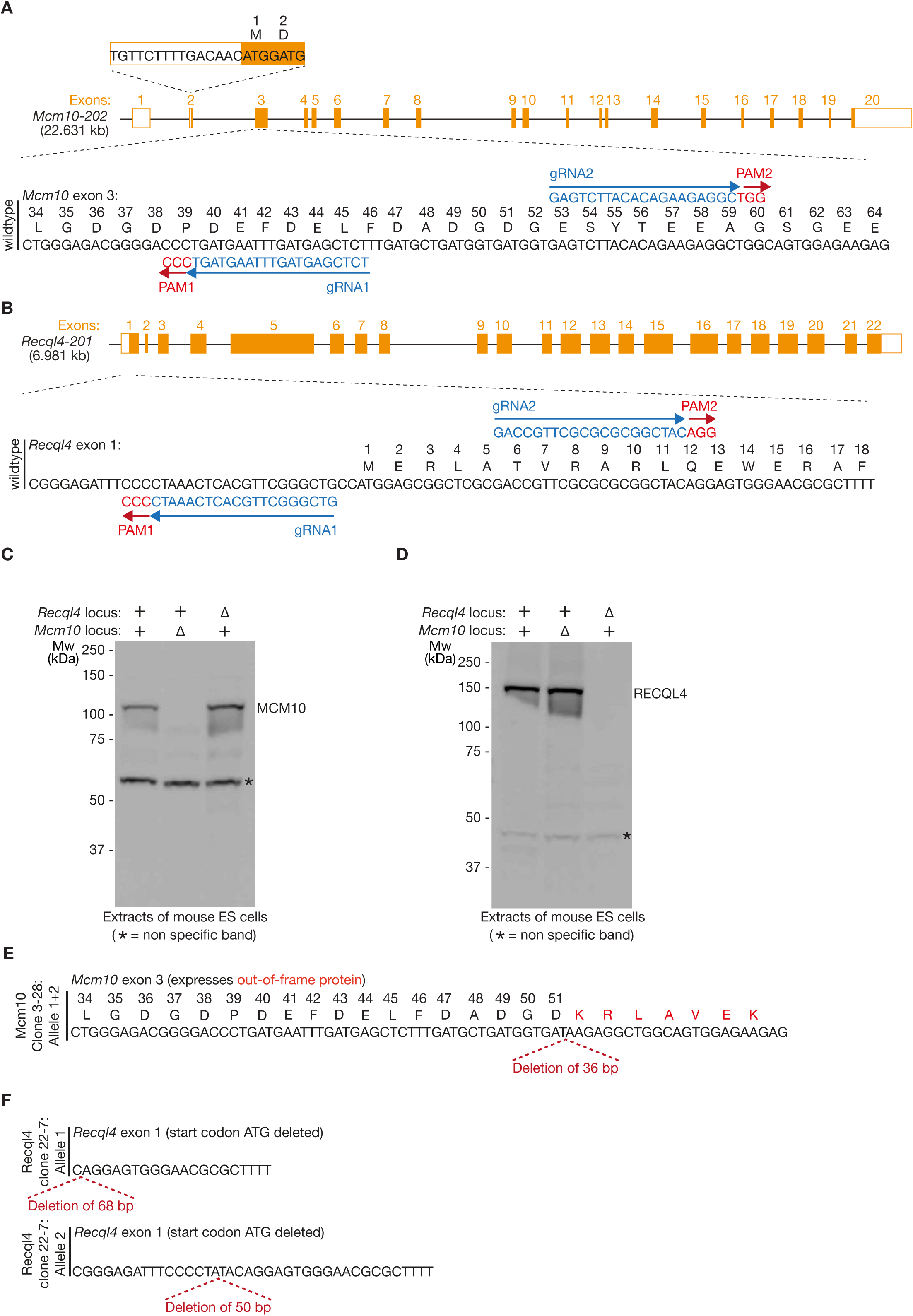
Small deletions at the start of the coding sequence of *Mcm10* or *Recql4* impair protein expression but do not cause inviability in mouse ES cells. (**A**) Depiction of the mouse *Mcm10* locus, showing the location of two guide RNAs (gRNAs) that were used in combination with Cas9-D10A to make small deletions in exon 3. PAM = Protospacer Adjacent Motif. (**B**) Equivalent illustration of the mouse *Recql4* locus, with corresponding gRNAs that were used to make small deletions in exon 1. (**C** - **D**) Clones generated with the gRNAs shown in (A-B) were analysed by immunoblotting. (**E**) DNA sequence analysis of the *Mcm10Δ* clone from (C). (**F**) DNA sequence analysis of the *Recql4Δ* clone from (D).

**Figure EV7.**
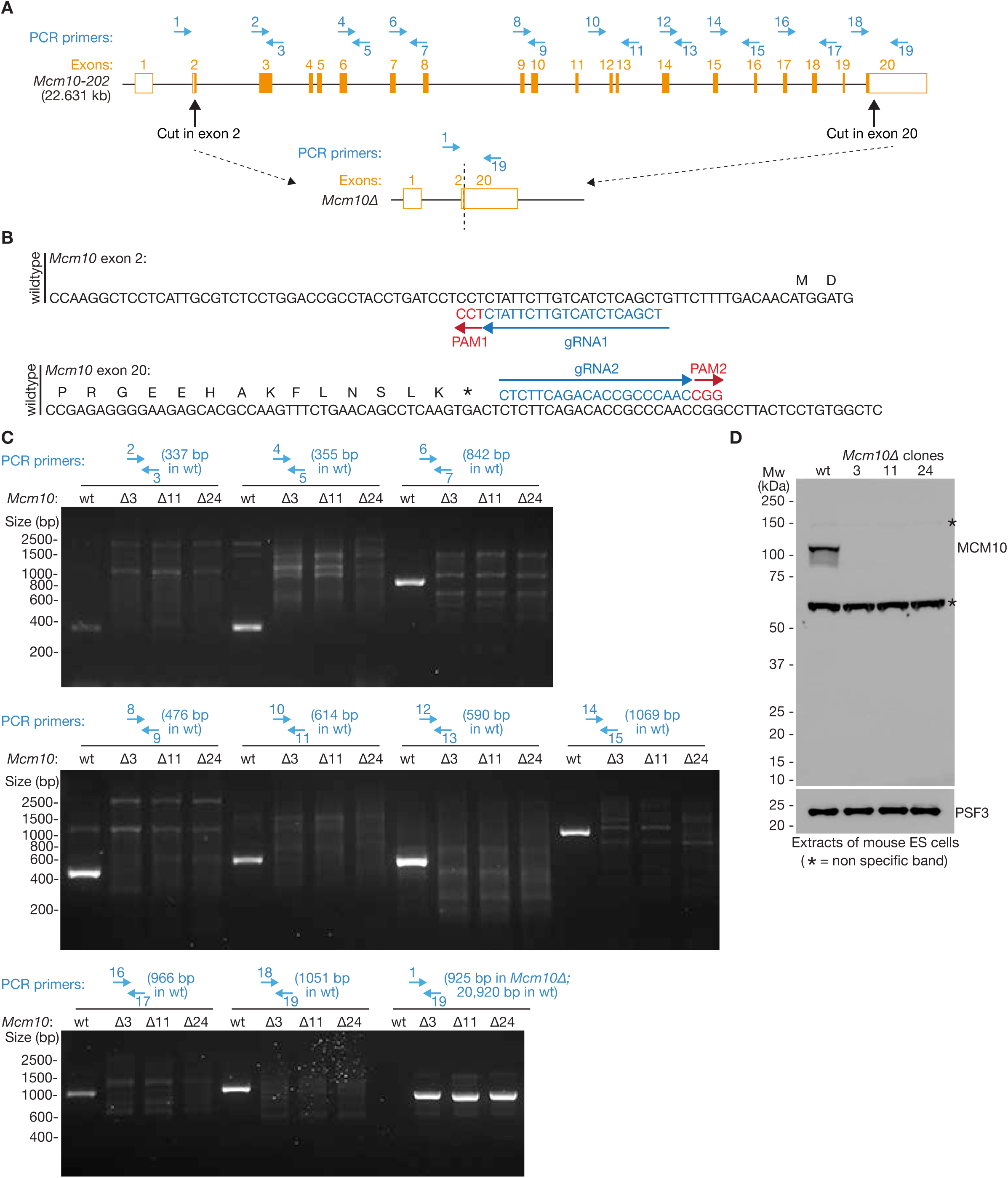
Complete deletion of the *Mcm10* gene is viable in mouse ES cells. (**A**) Depiction of the mouse *Mcm10* locus, showing the location of the sites in exon 2 and exon 20 where wild type Cas9 was directed by single gRNAs to cut both strands of the DNA template, leading to deletion of the entire gene (indicated at the bottom). Blue arrows indicate PCR primers that were used to monitor the locus. (**B**) DNA sequence of the relevant sections of exon 2 and exon 20 of Mcm10, showing each of the two guide RNAs (gRNAs) that were used in combination with wild type Cas9. PAM = Protospacer Adjacent Motif. (**C**) Genomic DNA from wild type mouse ES cells (wt) and three Mcm10Δ clones (Δ3, Δ11 and Δ24) was analysed by PCR with the indicated primer pairs. For each pair, the size of the expected band is shown in brackets. (**D**) Cell extracts of wild type and the three *Mcm10Δ* clones were analysed by immunoblotting.

**Figure EV8.**
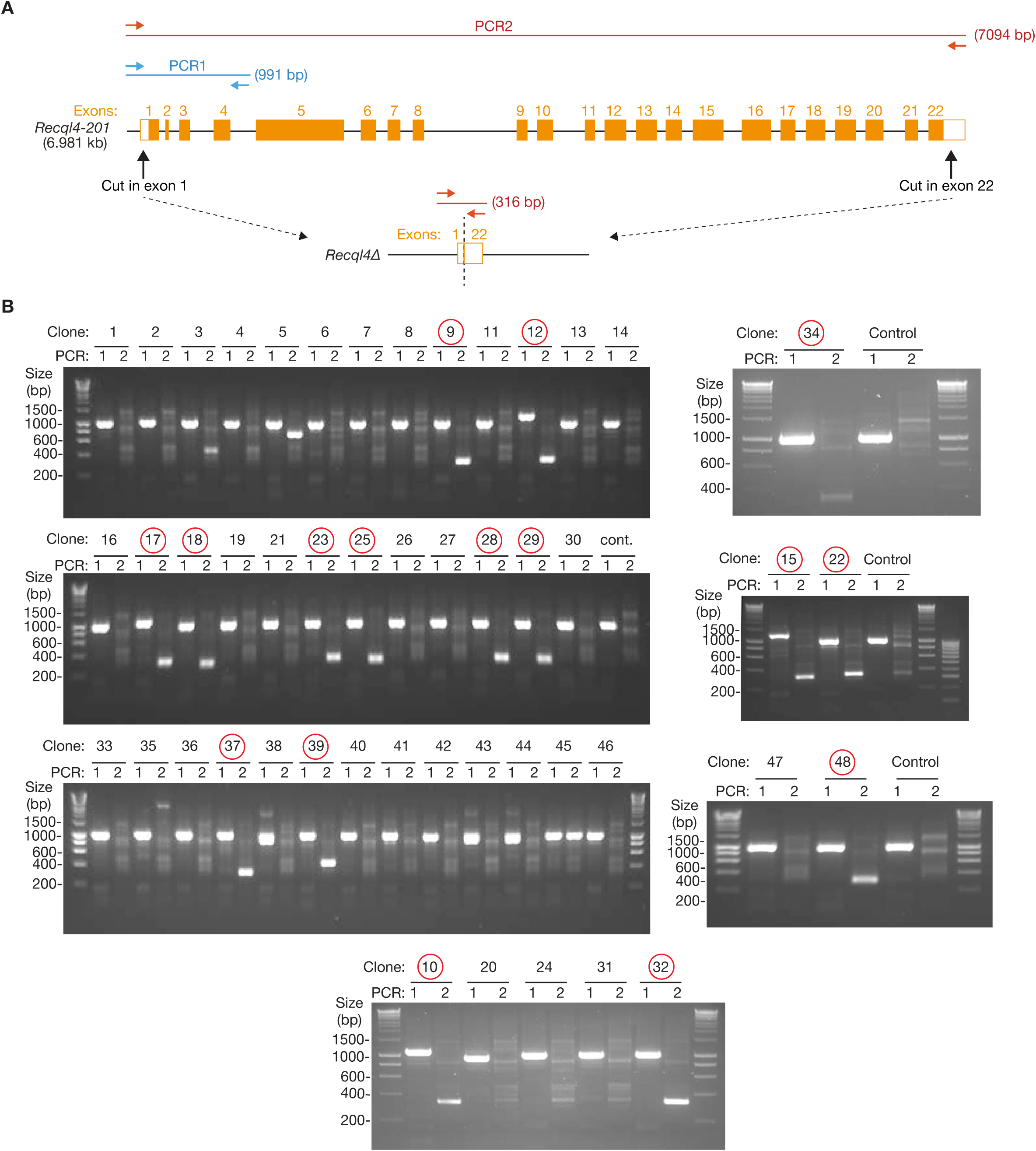
Complete deletion of the *Recql4* gene is only viable in heterozygous mouse ES cells. (**A**) Depiction of the mouse *Recql4* locus, showing the location of the sites before exon 1 and in exon 22 where wild type Cas9 was directed by single gRNAs to cut both strands of the DNA template, leading to deletion of the entire gene (indicated at the bottom). PCR1 (blue) produced a 991 bp product that indicates the presence of the wild type locus, whereas PCR2 (red) produced a 316 bp product that is indicative of complete deletion of the *Recql4* gene (amplification of the 7094 bp product for PCR2 from the wild type locus was too inefficient under these conditions and so was not observed). (**B**) Genomic DNA from wild type mouse ES cells (Control), and from 48 candidate Recql4Δ clones, was analysed using PCR1 and PCR2 that are summarised in (A). The sixteen clones shown in red were all heterozygous for the *Recql4* locus, with one allele producing a PCR2 product that indicates a complete deletion of the *Recql4* gene, and the second generating a PCR1 product of approximately wild type size.

**Figure EV9.**
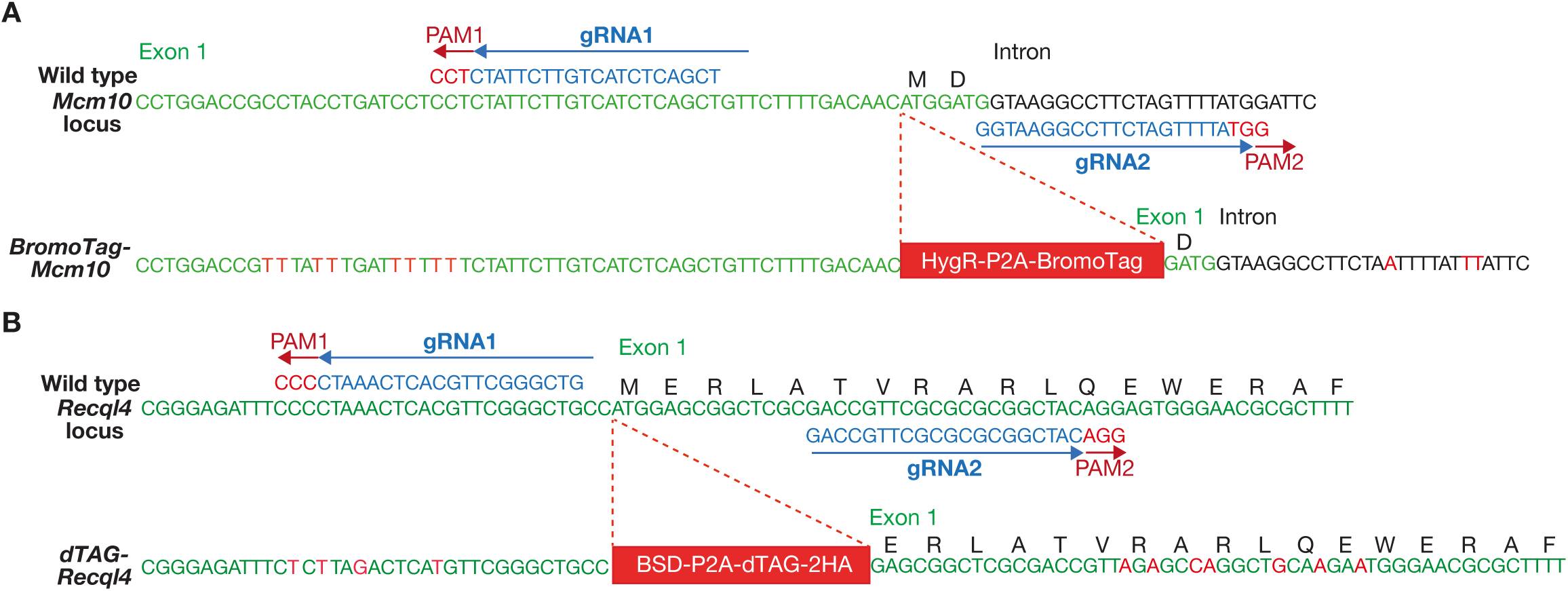
Generation of *BromoTag-Mcm10 dTAG-Recql4* mouse ES cells by genome editing with CRISPR-Cas9. (**A**) Exon 1 of the mouse *Mcm10* locus in wild type cells and *BromoTag-Mcm10*. Wild type cells were transfected with plasmids expressing gRNA1 and gRNA2 (PAM = Protospacer Adjacent Motif), together with the Cas9-D10A nickase, and a donor vector to allow repair of the locus after cutting, thereby inserting HygR-P2A-BromoTag (HygR = coding sequence for Hygromycin resistance gene; P2A = ribosome skipping sequence from porcine teschovirus-1 2A, which leads to expression of BromoTag-MCM10 as a separate protein to the HygR-P2A sequence). The 5’ and 3’ homology sequences in the donor vector also contained silent mutations (shown in red) that prevented re-cleavage by Cas9-D10A. (**B**) Exon 1 of the mouse *Recql4* locus in wild type cells and *dTAG-Recql4*. As above, the figure shows the location of the gRNAs that were used to direct Cas9-D10A to exon 1 of *Recql4*, allowing repair via a donor vector containing BSD-P2A-dTAG-2HA (BSD = coding sequence for Blastocidin resistance gene; P2A leads to expression of dTAG-RECQL4 as a separate protein to the BSD-P2A sequence; 2HA = two copies of the nine amino acid tag from human influenza hemagglutinin (HA) protein). As above, the 5’ and 3’ homology sequences in the donor vector also contained silent mutations (shown in red) that prevented re-cleavage by Cas9-D10A.

**Figure EV10.**
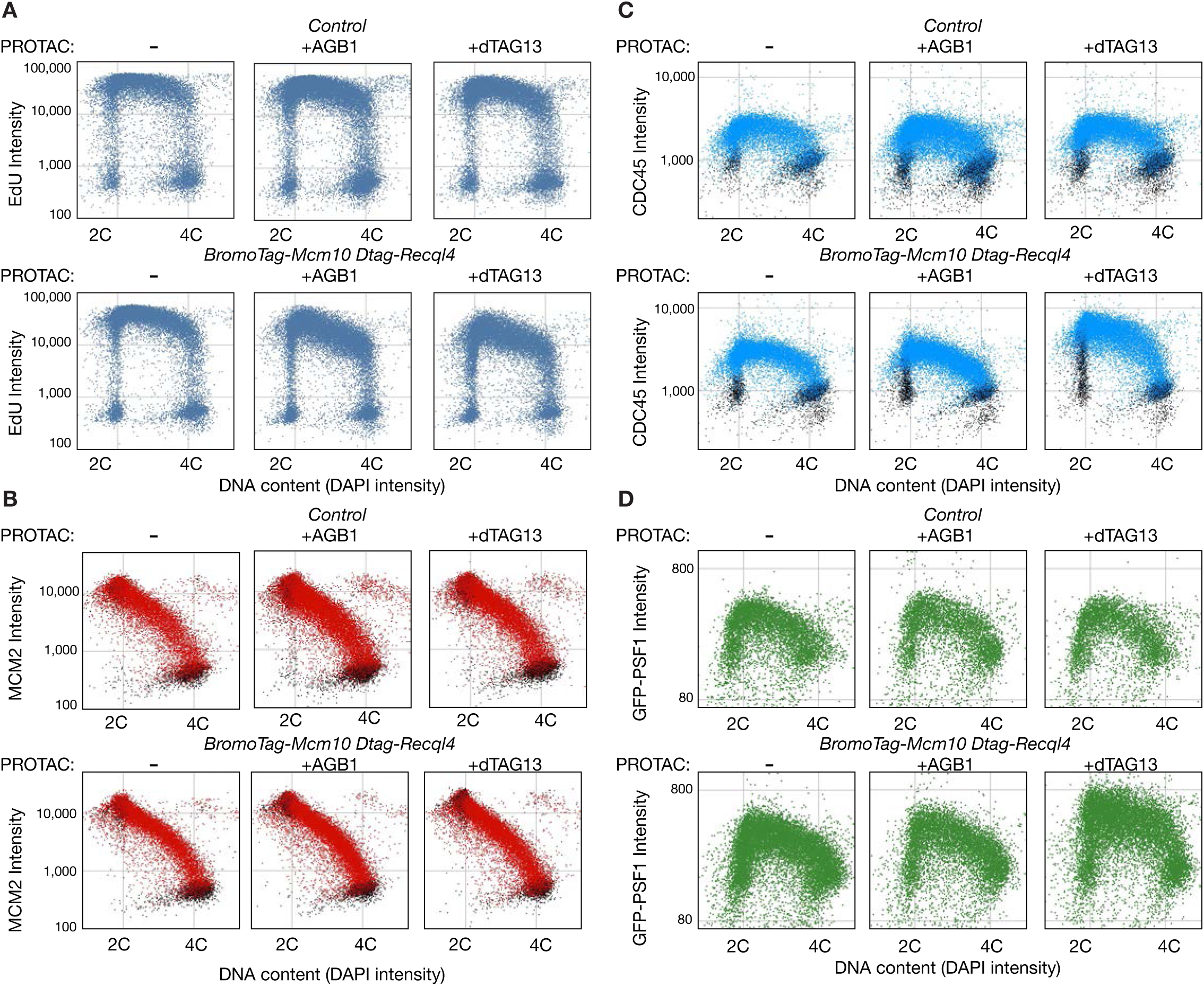
Depletion of RECQL4 has a greater impact on CMG helicase activation in mouse ES cells than depletion of MCM10. (**A**-**D**) correspond to data from the experiment in Figure 7F-J, for control and *BromoTag-Mcm10 dTAG-Recql4*, grown for eight hours as indicated in the absence of PROTACs, or in the presence of 250 nM AGB1, or 250 nM dTAG13. Cells were processed as described above for Figure 7F-J. For panels (B-C), EdU-negative cells (see Methods) are indicated in black. For panel (D), cells also expressed GFP-PSF1.

**Appendix Table S1:**
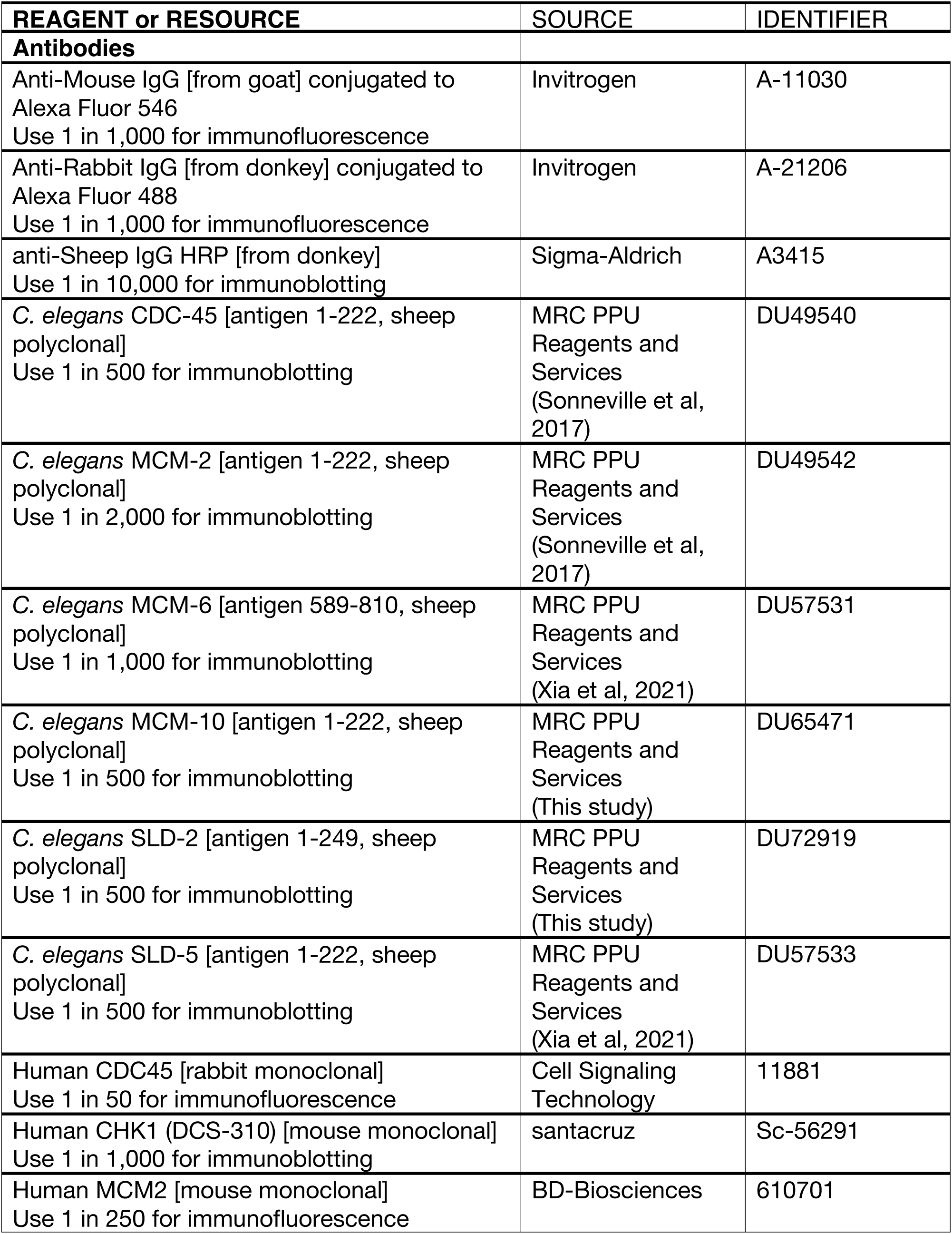
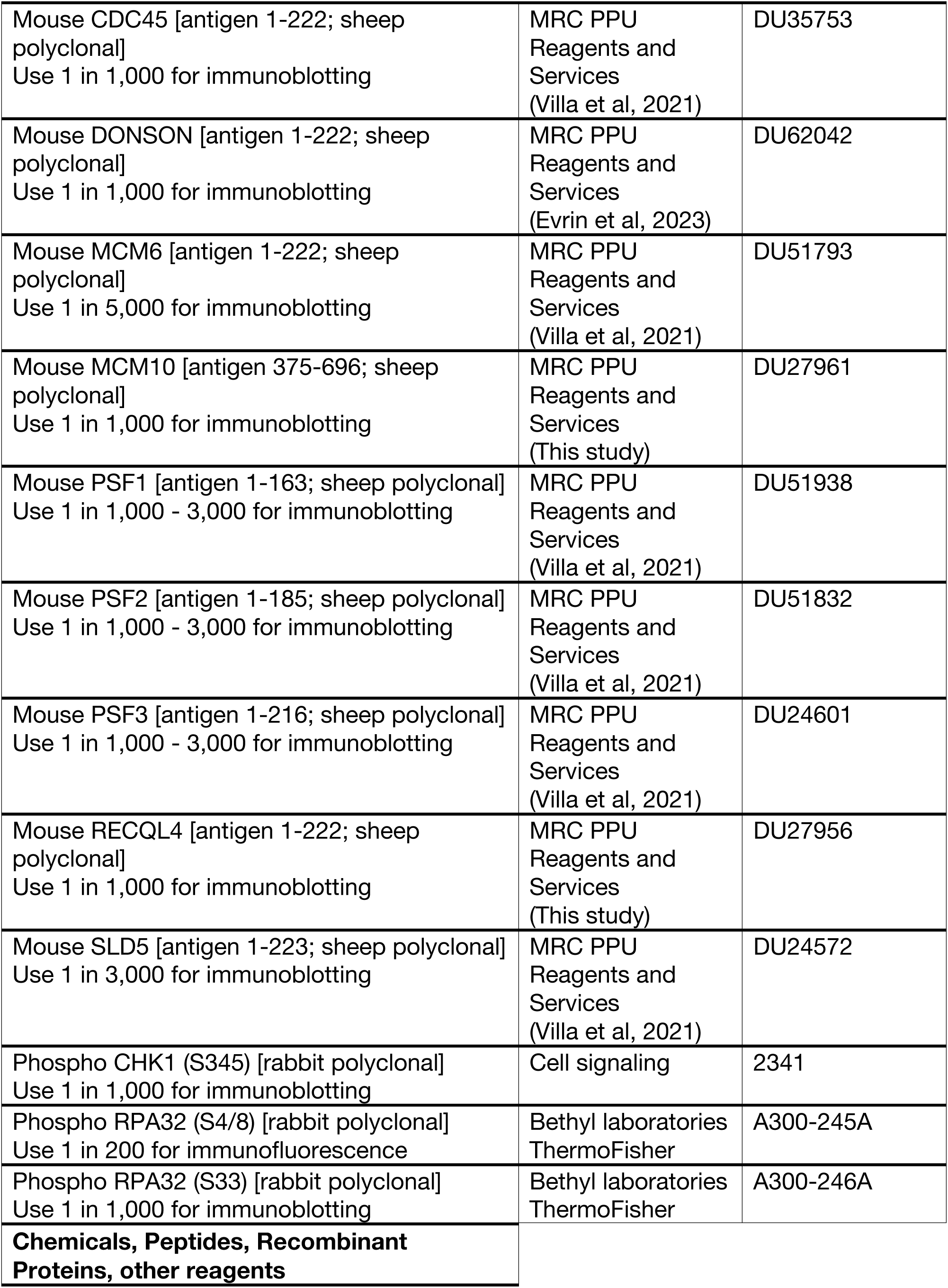
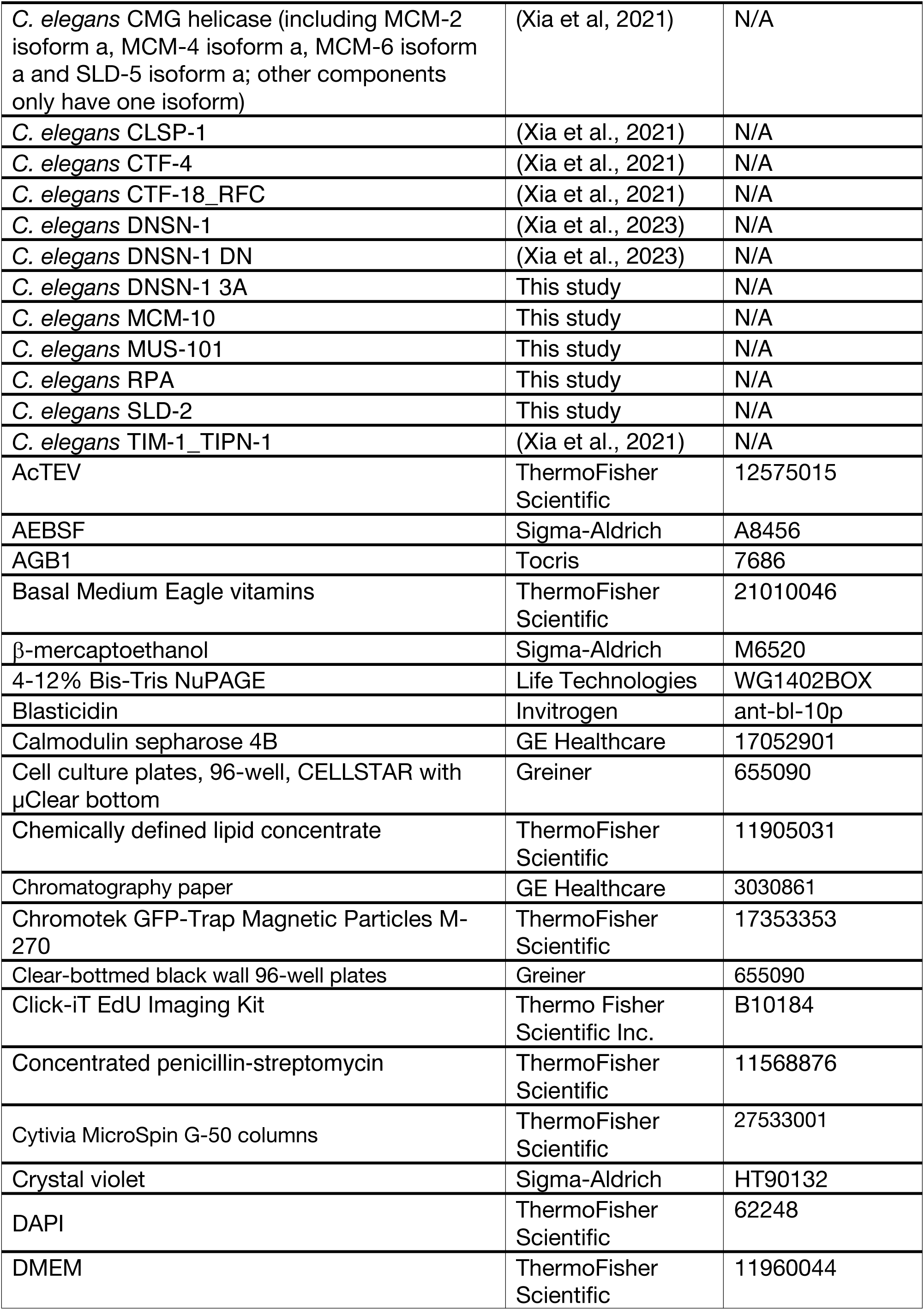
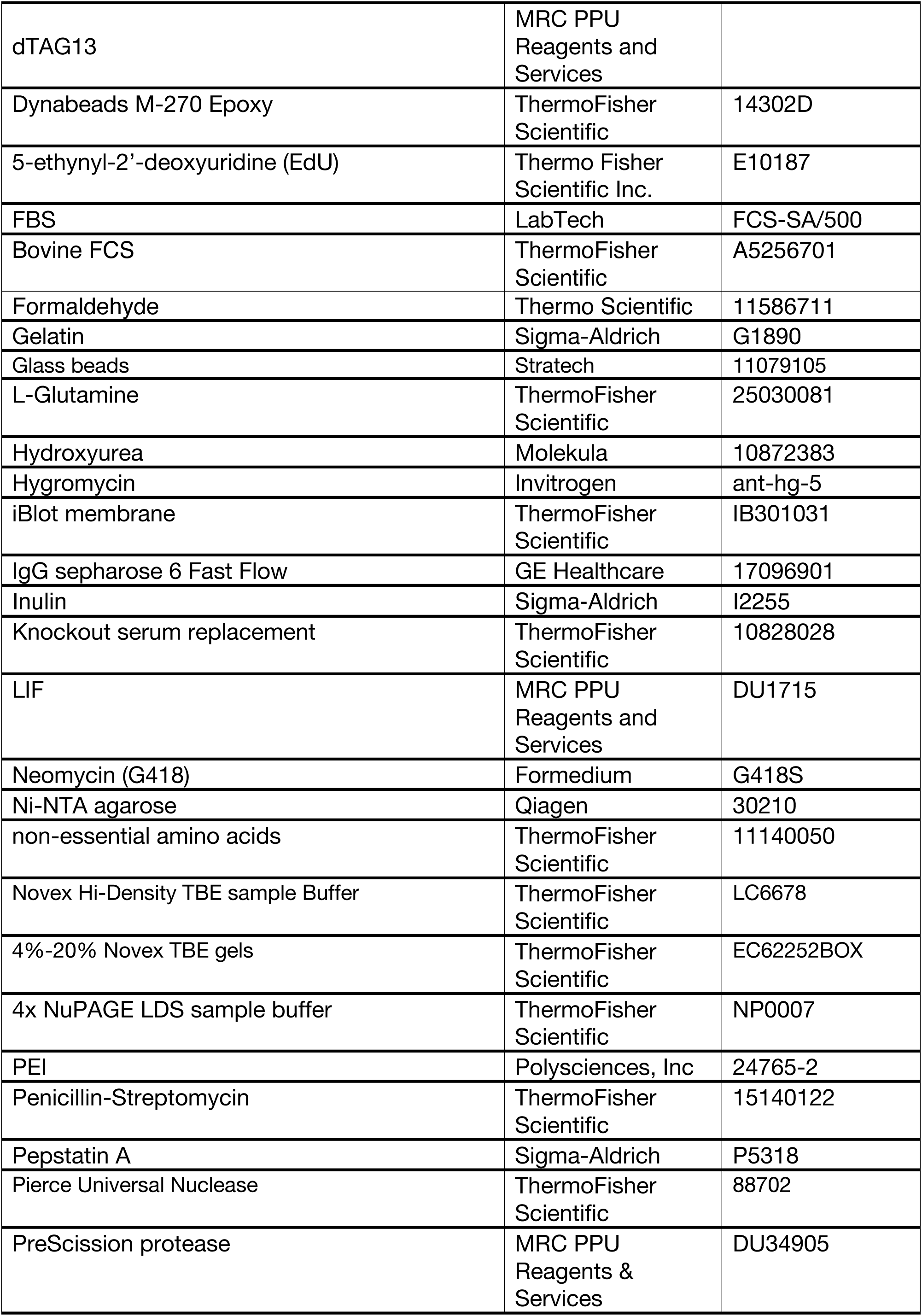
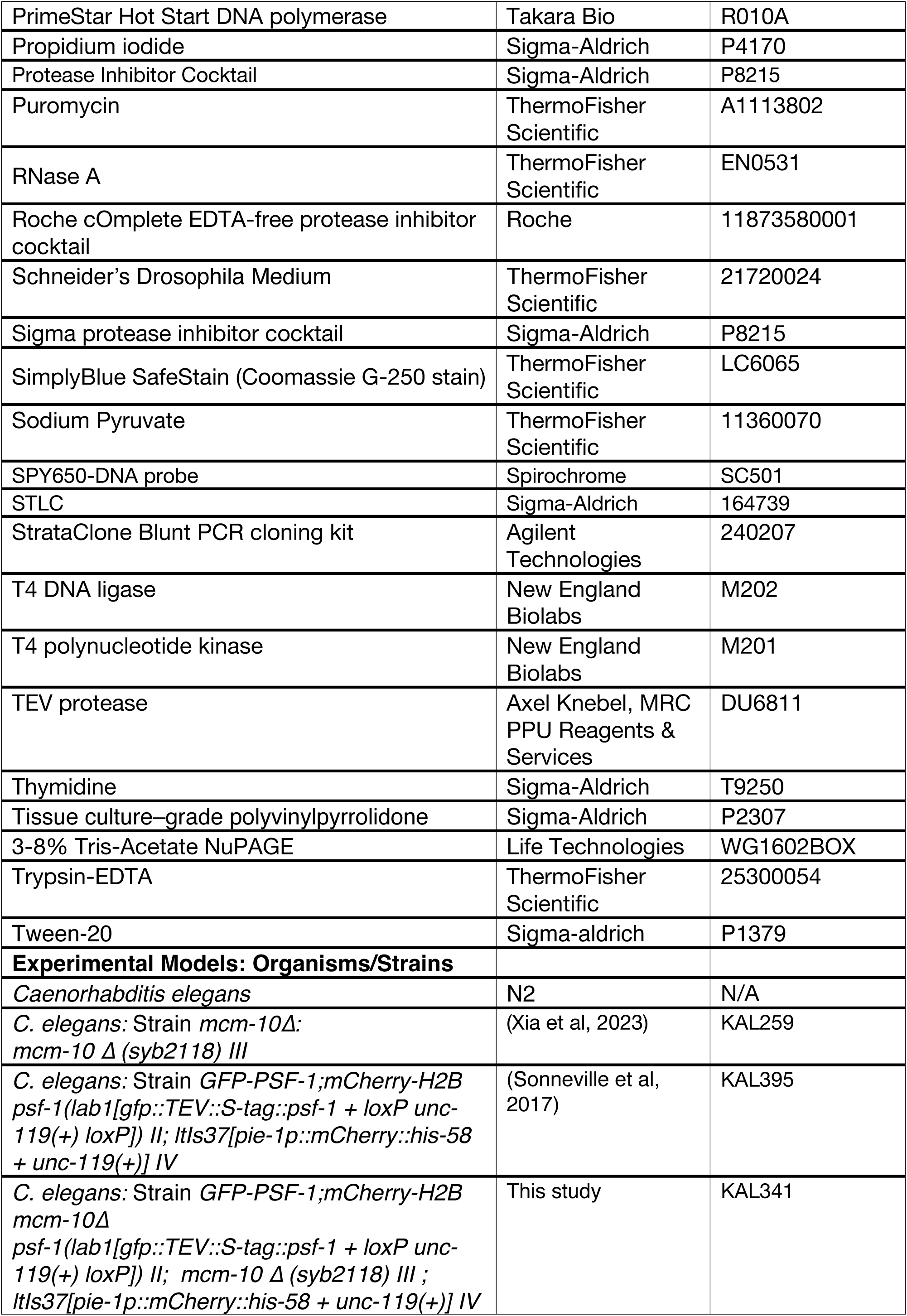
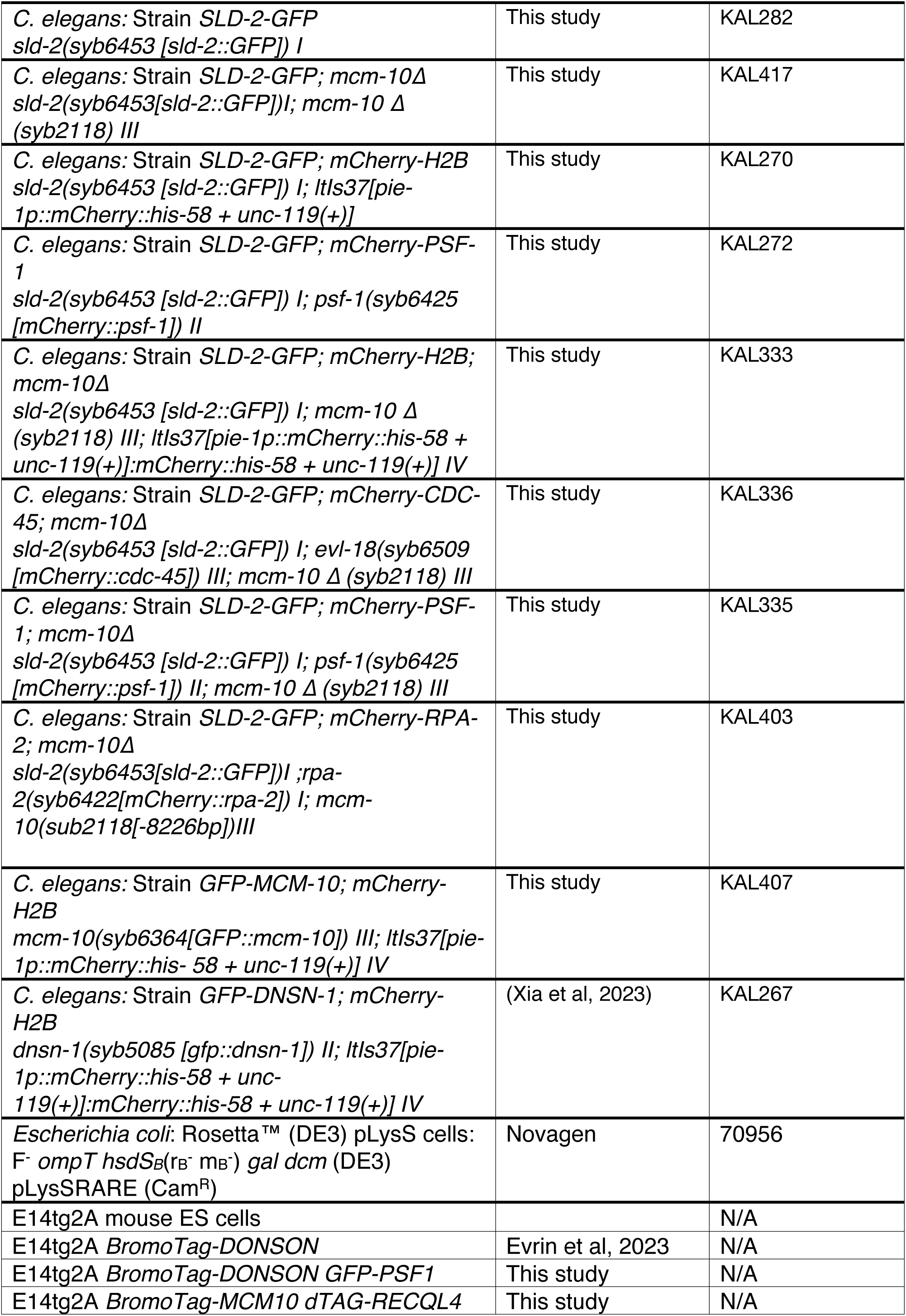
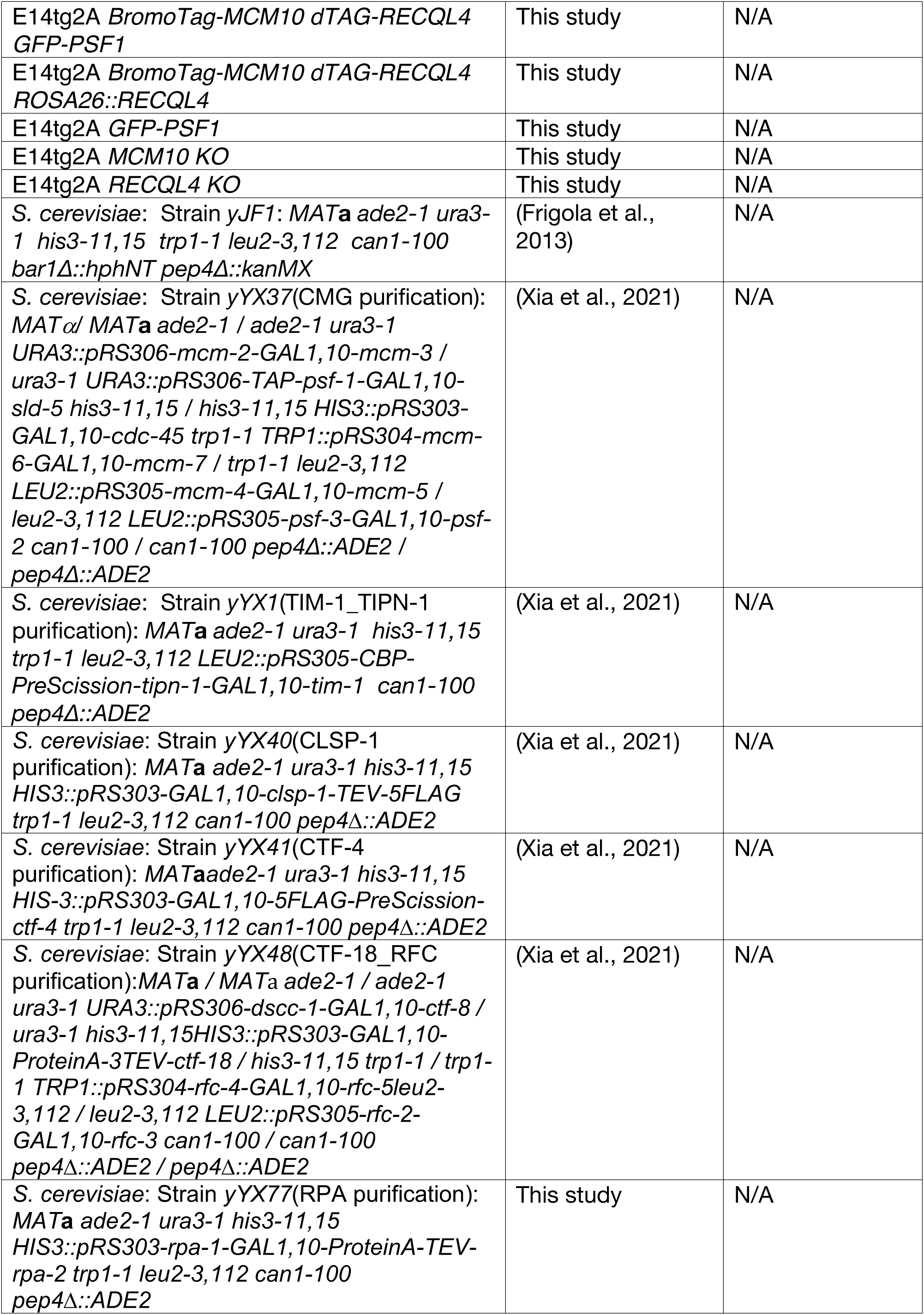
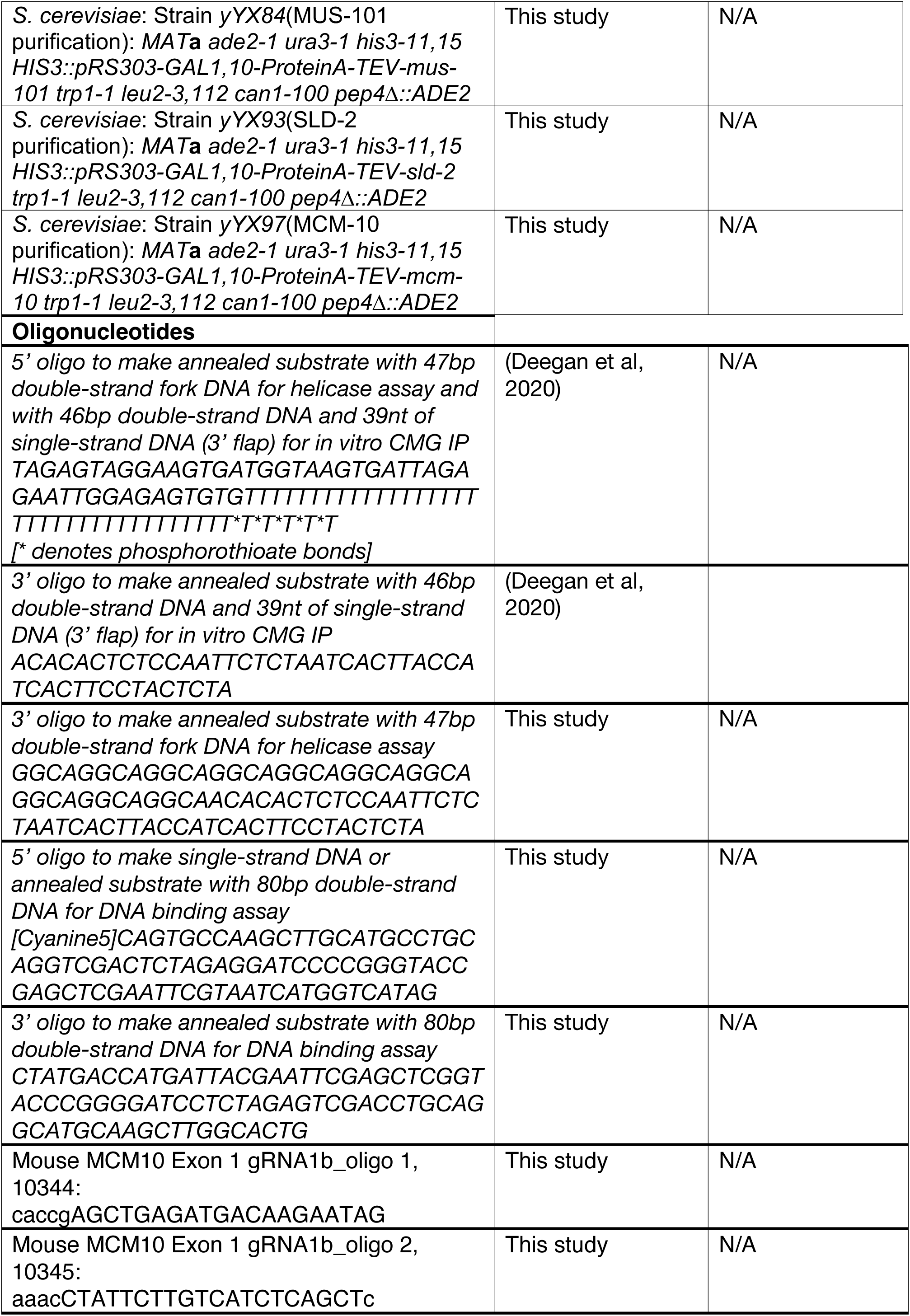
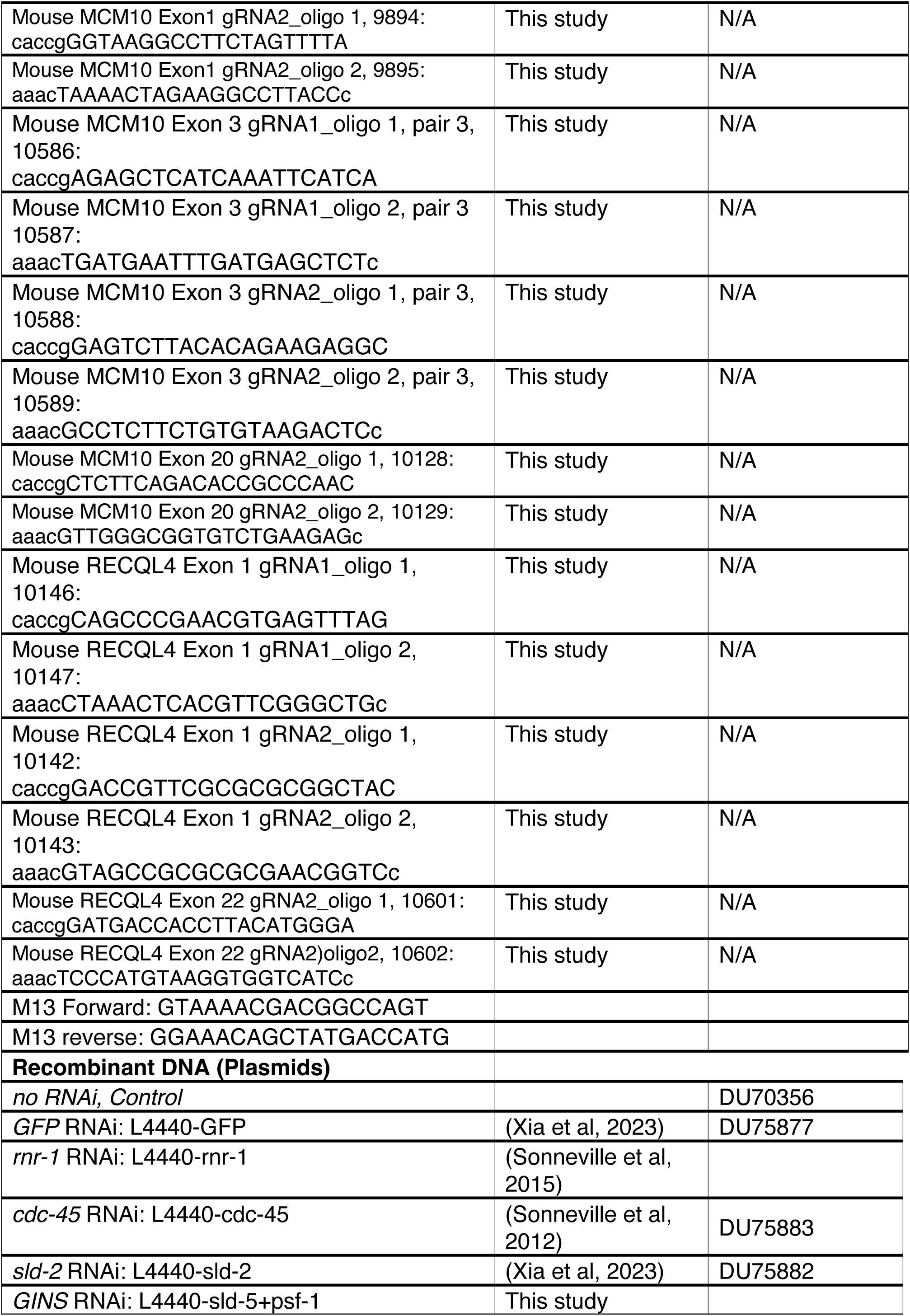
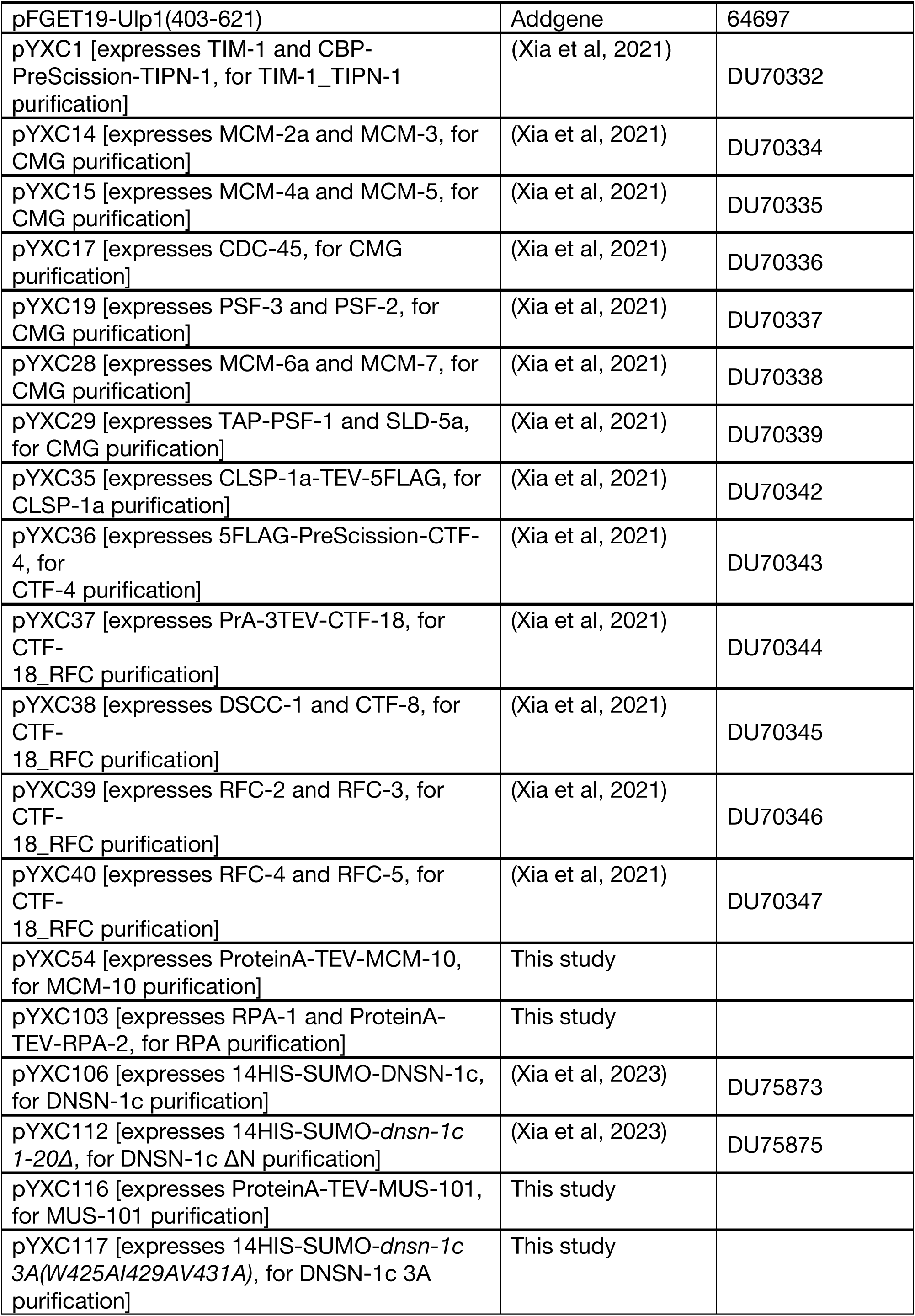
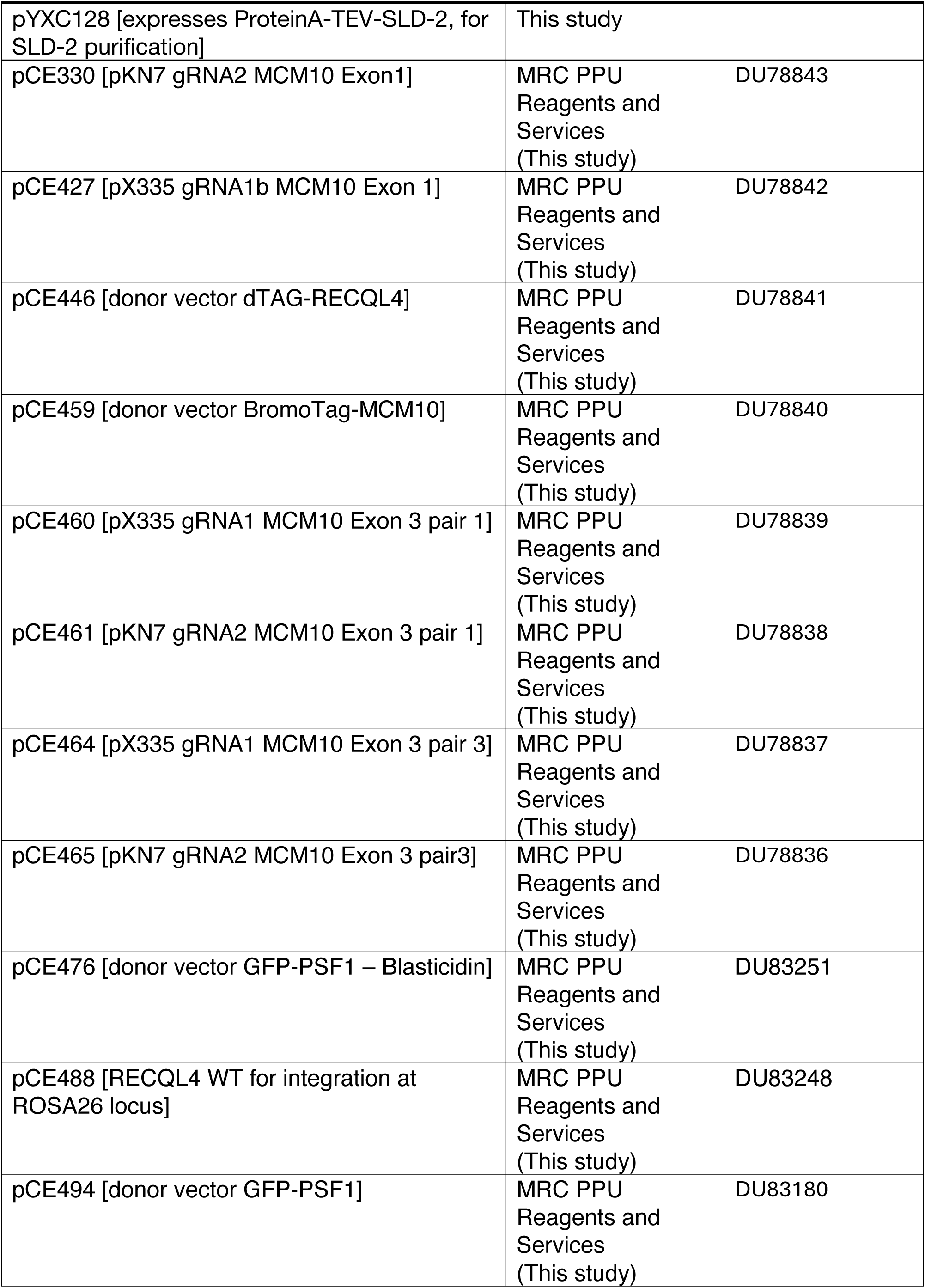
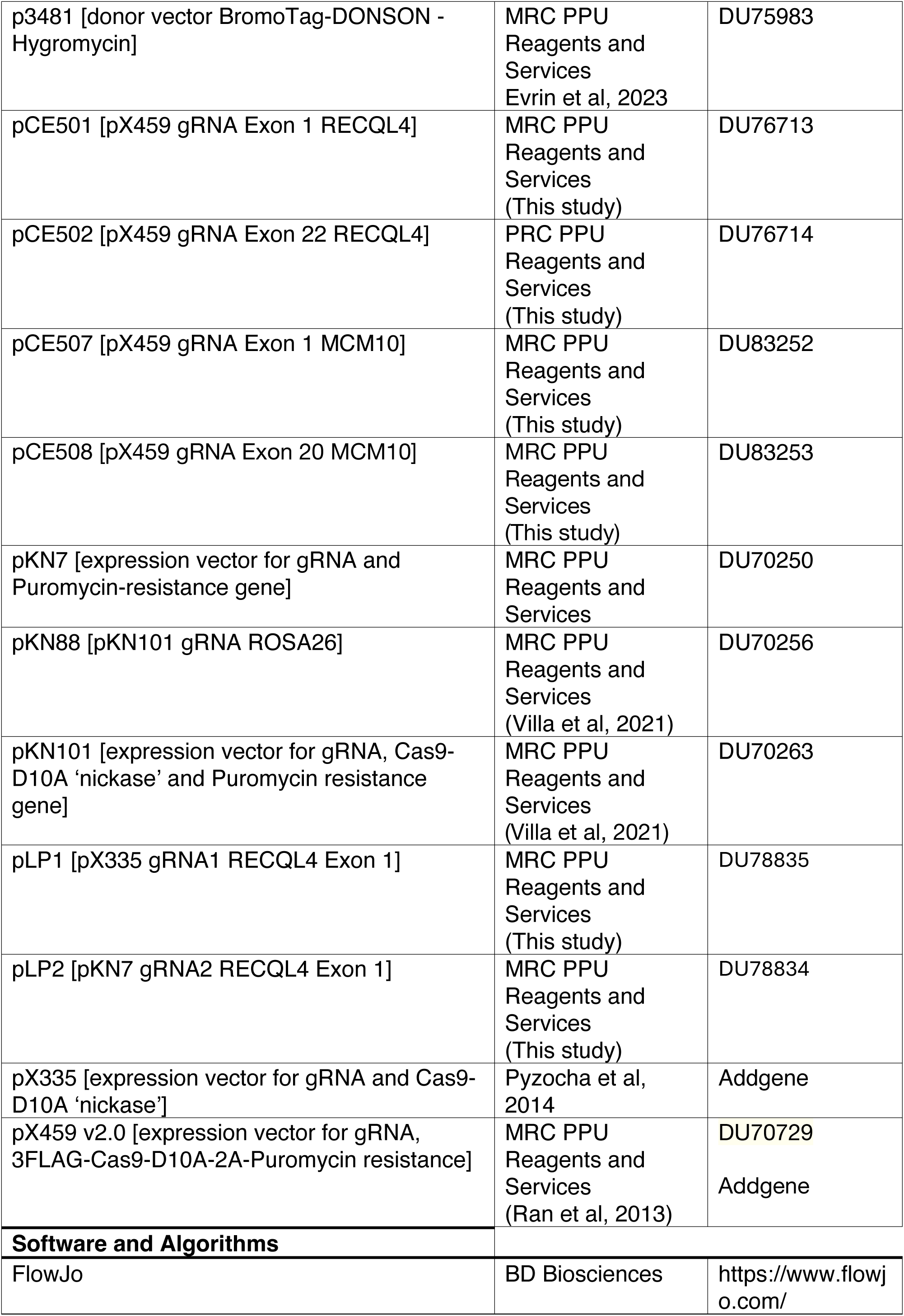
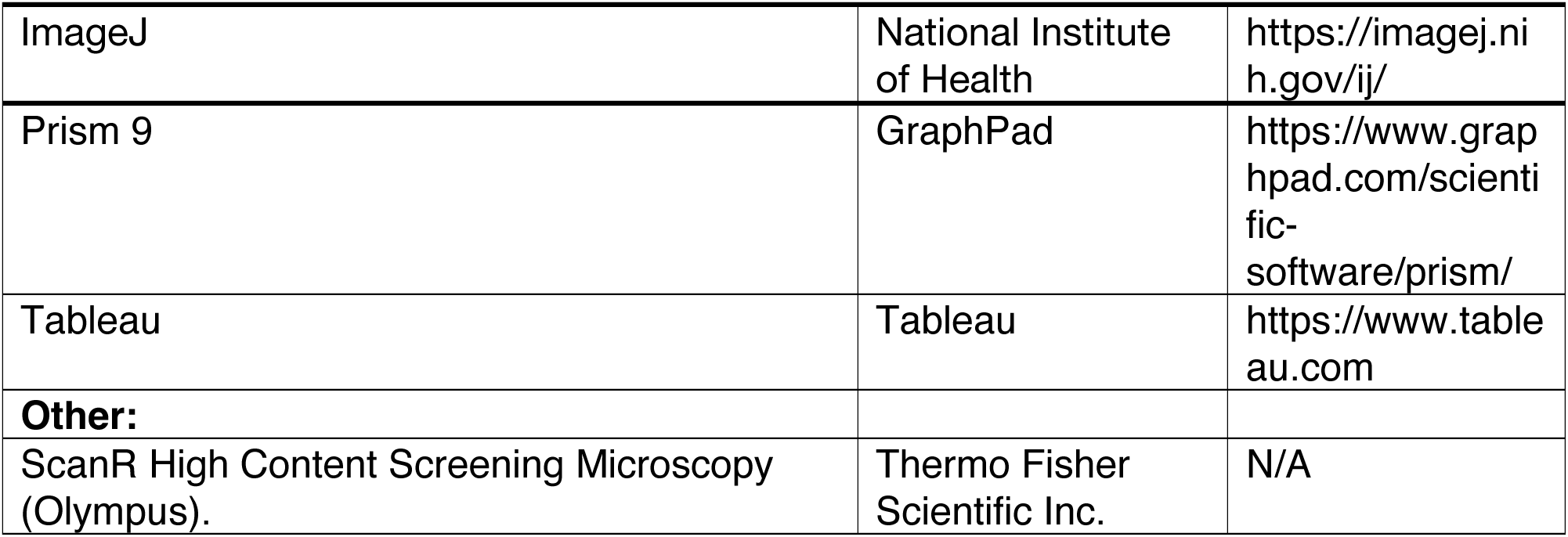
Reagents and resources used in this study.

